# Genetic determinants of *EGFR*-Driven Lung Cancer Growth and Therapeutic Response *In Vivo*

**DOI:** 10.1101/2020.04.13.036921

**Authors:** Giorgia Foggetti, Chuan Li, Hongchen Cai, Jessica A. Hellyer, Wen-Yang Lin, Deborah Ayeni, Katherine Hastings, Jungmin Choi, Anna Wurtz, Laura Andrejka, Dylan Maghini, Nicholas Rashleigh, Stellar Levy, Robert Homer, Scott Gettinger, Maximilian Diehn, Heather A. Wakelee, Dmitri A. Petrov, Monte M. Winslow, Katerina Politi

**Affiliations:** Yale Cancer Center, Yale School of Medicine, New Haven, Connecticut; Department of Biology, Stanford University School of Medicine, Stanford, California; Department of Genetics, Stanford University School of Medicine, Stanford, California; Stanford Cancer Institute, Department of Medicine, Division of Oncology, Stanford University School of Medicine, Stanford, California; Department of Genetics, Yale School of Medicine, New Haven, Connecticut; Department of Biomedical Sciences, Korea University College of Medicine, Seoul, Korea; Department of Pathology, Yale School of Medicine, New Haven, Connecticut; VA Connecticut HealthCare System, Pathology and Laboratory Medicine Service, West Haven, Connecticut; Department of Medicine (Section of Medical Oncology), Yale School of Medicine, New Haven, Connecticut; Institute for Stem Cell Biology and Regenerative Medicine, Stanford University School of Medicine, Stanford, California; Department of Radiation Oncology, Stanford University School of Medicine, Stanford, California; Department of Pathology, Stanford University School of Medicine, Stanford, California; Cancer Biology Program, Stanford University School of Medicine, Stanford, California

## Abstract

Cancer genome sequencing has uncovered substantial complexity in the mutational landscape of tumors. Given this complexity, experimental approaches are necessary to establish the impact of combinations of genetic alterations on tumor biology and to uncover genotype-dependent effects on drug sensitivity. In lung adenocarcinoma, *EGFR* mutations co-occur with many putative tumor suppressor gene alterations, however the extent to which these alterations contribute to tumor growth and their response to therapy *in vivo* has not been explored experimentally. By integrating a novel mouse model of oncogenic *EGFR*-driven *Trp53*-deficient lung adenocarcinoma with multiplexed CRISPR–Cas9-mediated genome editing and tumor barcode sequencing, we quantified the effects of inactivation of ten putative tumor suppressor genes. Inactivation of *Apc*, *Rb1*, or *Rbm10* most strongly promoted tumor growth. Unexpectedly, inactivation of *Lkb1* or *Setd2 –* which are the strongest drivers of tumor growth in an oncogenic *Kras*-driven model – reduced *EGFR*-driven tumor growth. These results are consistent with the relative frequency of these tumor suppressor gene alterations in human *EGFR-* and *KRAS*-driven lung adenocarcinomas. Furthermore, *Keap1* inactivation reduces the sensitivity of *EGFR*-driven *Trp53*-deficient tumors to the EGFR inhibitor osimertinib. Importantly, in human *EGFR/TP53* mutant lung adenocarcinomas, mutations in the KEAP1 pathway correlated with decreased time on tyrosine kinase inhibitor treatment. Our study highlights how genetic alterations can have dramatically different biological consequences depending on the oncogenic context and that the fitness landscape can shift upon drug treatment.

---

During tumor evolution, cancer cells accumulate alterations in oncogenes and tumor suppressor genes, which contribute to many of the hallmarks of cancer^1^. Despite their extensive genomic complexity, tumors are frequently classified based on the presence of a single oncogenic driver mutation, while the function of co-incident tumor suppressor gene alterations is largely ignored. There is emerging evidence that the interplay between oncogenic drivers and tumor suppressor genes may influence tumor fitness and impact treatment response^2^. However, the combinatorially vast landscape of genomic alterations makes it difficult, except in the most extreme cases, to glean information about the epistatic interactions between tumor suppressor genes and oncogenes from human cancer sequencing data alone^3^. This complexity makes inferring the relationship between genotype and therapy response even more tenuous.

Recently, high-throughput, tractable systems that combine autochthonous mouse modeling and genome editing have been developed to directly uncover the functional consequences of genetic alterations during tumorigenesis *in vivo*^4–10^. However, very few studies have investigated the biological consequences of inactivating tumor suppressor genes in the context of different oncogenic drivers *in vivo*, and existing knowledge about the role of specific genetic alterations in tumor suppressor genes has been primarily inferred from correlative human studies^11–13^.

In lung adenocarcinoma, *EGFR* and *KRAS* are the two most frequently mutated oncogenic driver genes and occur within a background of diverse putative tumor suppressor gene alterations^2,11,12,14^. Among these, *TP53* is the most commonly mutated tumor suppressor gene in both oncogenic *EGFR-* and *KRAS*-driven lung adenocarcinoma, consistent with the importance of disrupting this pathway during lung cancer development^2,13,15–18^. Interestingly, many other putative tumor suppressor genes are mutated at different frequencies in oncogenic *EGFR-* and *KRAS-*driven lung adenocarcinomas^2,11,15^. Whether these differences are due to different fitness effects that depend on the oncogenic context has never been tested experimentally. Indeed, previous studies on tumor suppressor genes in lung cancer models *in vivo* have been primarily conducted in the context of oncogenic *Kras*-driven tumors, while the functional importance of different tumor suppressor genes in *EGFR*-driven lung tumors remains largely unstudied (**Supplemental Fig. 1a**).

In addition to driving growth, inactivation of tumor suppressor pathways may affect the therapeutic response to therapies. In advanced *EGFR* mutant lung adenocarcinomas, treatment with EGFR tyrosine kinase inhibitors (TKIs) is the first-line of therapy^19–22^. Response rates to TKIs are high, however, there is large variability in the depth and duration of response between patients, and acquired resistance inevitably occurs^14^. Genomic alterations, including those in *RB1* and *TP53*, have been found to correlate with clinical responses to TKIs^13,17,23^. However, given the complexity and diversity of genomic alterations in these tumors, the functional contribution of individual genes to drug resistance remains poorly understood.

To quantify the functional importance of a panel of ten diverse putative tumor suppressor genes on oncogenic *EGFR-*driven lung tumor growth *in vivo,* we coupled multiplexed CRISPR–Cas9-mediated somatic genome editing and tumor barcoding sequencing (Tuba-seq) with a novel genetically engineered mouse model of *EGFR*^*L858R*^-driven *Trp53-*deficient lung cancer. Through the comparison of tumor suppressor effects in oncogenic *EGFR-* and *Kras*-driven lung cancer models, we uncovered prevalent epistasis between the oncogenic drivers and tumor suppressor genes, which explains the different mutational spectra of these tumor suppressor genes in oncogenic *EGFR*- and *KRAS*-driven human lung adenocarcinomas. Moreover, we established a direct causal link between tumor suppressor genotypes and differential responses to osimertinib treatment in *EGFR*-driven lung adenocarcinoma that are supported by correlative human mutational datasets.

## Results

### Development of a lentiviral-Cre based mouse model of oncogenic *EGFR*-driven lung adenocarcinoma

The use of lentiviral vectors to initiate tumors in genetically engineered mouse models of human cancer enables control of tumor multiplicity, tumor barcoding to map clonality, and the delivery of lentivirus-encoded cDNAs, shRNAs, and sgRNAs to modify neoplastic cells^6,8,24,25^. The simplicity of viral-Cre initiated models of oncogenic *Kras*-driven lung cancer has enabled the analysis of many genes that co-operate to drive tumor growth within these autochthonous mouse models^26^ (**Supplemental Fig. 1a**). To permit the generation of virally-initiated oncogenic *EGFR*-driven *Trp53*-deficient lung tumors, we bred mice with an rtTA/tetracycline-inducible transgene encoding the common lung adenocarcinoma-associated EGFR^L858R^ mutant (*TetO-EGFR*^*L858R*^), a Cre-regulated *rtTA* transgene (*Rosa26*^*CAGs-LSL-rtTA3-IRES-mKate*^, abbreviated *R26*^*RIK*^), and homozygous *Trp53* floxed alleles (*p53*^*flox/flox*^)^27–29^. In these *TetO-EGFR*^*L858R*^;*Rosa26*^*RIK*^;*p53*^*flox/flox*^ (*EGFR;p53*) mice, lentiviral-Cre transduction of lung epithelial cells leads to the expression of rtTA and mKate as well as the inactivation of *Trp53*. Co-incident doxycycline treatment induces the expression of oncogenic EGFR (**Fig. 1a**).

**Fig. 1.**
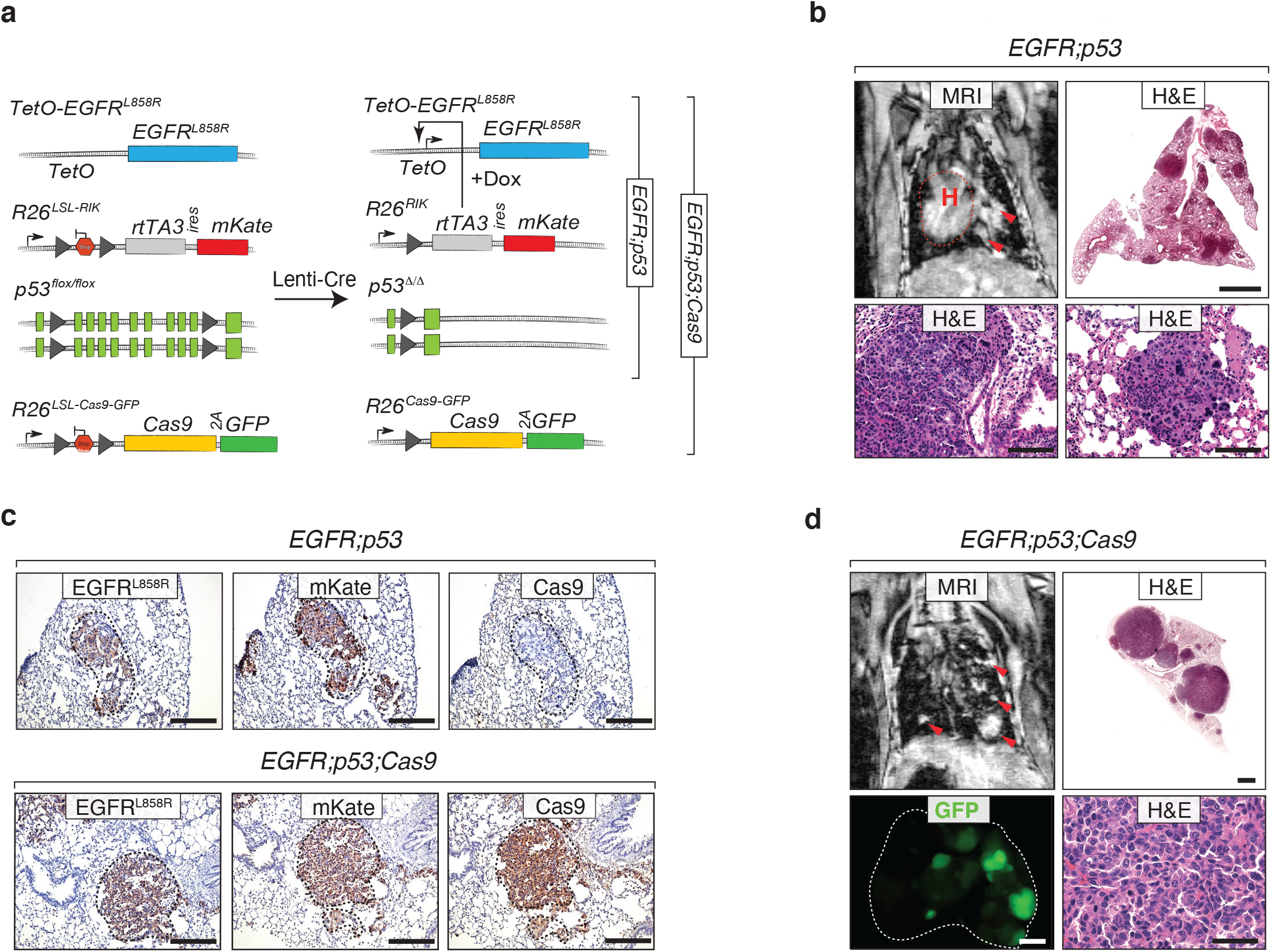
Lentiviral Cre-mediated lung tumor initiation in *EGFR;p53* and *EGFR;p53;Cas9* mice. **a**, Schematic of the *TetO-EGFR*^*L858R*^, *R26*^*LSL-RIK*^, and *p53*^*flox*^ alleles in *EGFR;p53* mice, prior to and following Cre-mediated recombination. Conditional expression of oncogenic EGFR^L858R^ is under the control of a tetracycline response element (*TetO*), which is induced by rtTA in the presence of doxycycline (Dox). Lentiviral-Cre inactivates *Trp53* and enables the expression of the reverse tetracycline-regulated transactivator (*rtTA3*) and mKate (from the *R26*^*LSL-RIK*^ allele). Cre also allows expression of Cas9 and GFP (from the *R26*^*LSL-Cas9-GFP*^ allele) in *EGFR;p53;Cas9* mice. **b**, MRI showing tumor development 11 weeks after tumor initiation in *EGFR;p53* mice (top left panel) with 2×10^6^ ifu Lenti-Cre. The dashed red line surrounds the heart (H) and red arrows indicate areas of tumor. Hematoxylin and eosin (H&E) staining shows lung adenocarcinoma development. Scale bars = 1.2 mm and 100 μm in the top and bottom panels, respectively. Images are from a representative mouse (*N* = 5). **c**, Immunostaining for EGFR^L858R^, mKate and Cas9 in tumors in *EGFR;p53* and *EGFR;p53;Cas9* mice. The dashed lines indicate areas of tumor. Scale bars = 200 μm. **d**, MRI and H&E showing tumor development 16 weeks after tumor initiation in *EGFR;p53;Cas9* mice with 1×10^5^ ifu Lenti-Cre.Tumors are positive for GFP in *EGFR;p53;Cas9* mice and lungs are indicated by the white dashed line. H&E image scale bars = 1.2 mm and 100 μm in top and bottom right panels, respectively. GFP image scale bar = 2.5 mm.

We initiated tumors with a lentiviral-PGK-Cre vector^30^ in *EGFR;p53* mice and used magnetic resonance imaging (MRI) to monitor tumor development (**Fig. 1b**). Tumors were first detectable in *EGFR;p53* mice 8 weeks after tumor initiation. Histological analysis of lungs 11 weeks after tumor initiation confirmed the development of multifocal lung adenocarcinomas (**Fig. 1b**, bottom panels). Tumors stained positively for EGFR^L858R^ and mKate, as well as surfactant protein C (SP-C) and the lung lineage-defining transcription factor NKX2.1/TTF-1 (**Fig. 1c**, top panels)^31–33^. Importantly, in this model, tumors were more focal than the diffuse tumors that rapidly develop in the previous *CCSP-rtTA;TetO-EGFR*^*L858R*^ model^27^ likely due to tumor initiation from fewer cells in the virally initiated model (**Fig. 1b**, top right panel). Lentivirus-induced tumors in *EGFR;p53* mice were more poorly differentiated than those typically observed in the *CCSP-rtTA;TetO-EGFR*^*L858R*^ model (**Fig. 1b**, bottom panels) as shown by increased prevalence of a micropapillary pattern, which is known to be highly aggressive in human adenocarcinoma^27^. Thus, this new lentiviral-Cre-initiated model recapitulates the genetic and histopathological features of human oncogenic *EGFR-*driven *TP53*-deficient lung tumors.

### Multiplexed quantification of tumor suppressor gene function in *EGFR*-driven lung tumors

To enable somatic genome editing in the *EGFR;p53* model, we further incorporated a conditional *Cas9* allele (*Rosa26*^*LSL-Cas9-GFP*^)^4^ to generate *TetO-EGFR*^*L858R*^;*R26*^*RIK/LSL-Cas9-GFP*^;*p53*^*flox/flox*^ (*EGFR;p53;Cas9*) mice (**Fig. 1a**). Lentiviral-Cre delivery to *EGFR;p53;Cas9* mice initiated multifocal lung adenomas and adenocarcinomas that expressed EGFR^L858R^, mKate, Cas9, and GFP (**Figs. 1c, d**). Tumors in *EGFR;p53;Cas9* mice were histologically similar to those in the *EGFR;p53* mice (**Figs. 1b, d** and **Supplemental Fig. 1b**).

We used an improved version of our Tuba-seq approach to quantify tumor suppressor gene function in oncogenic *EGFR*-driven lung tumors (**Methods**). Genomic integration of barcoded lentiviral vectors uniquely tags each transduced cell and all of the neoplastic cells within the resulting clonal tumors^8^. Each barcode encodes an 8-nucleotide sgID region specific to the sgRNA followed by a random 15-nucleotide barcode; thus, high-throughput sequencing of this sgID-BC region from bulk tumor-bearing lung can be used to quantify the number of cells in each tumor of each genotype (**Methods**)^8^. With this approach, the absolute number of neoplastic cells in each tumor is calculated by normalizing the number of reads of each unique barcode (sgID-BC) to the number of reads from benchmark control cells added to each sample (**Fig. 2a**, **Supplemental Fig. 9a and Methods**). Thus, Tuba-Seq enables the parallel analysis of the impact of multiple tumor suppressor gene alterations on tumor growth *in vivo.*

**Fig. 2.**
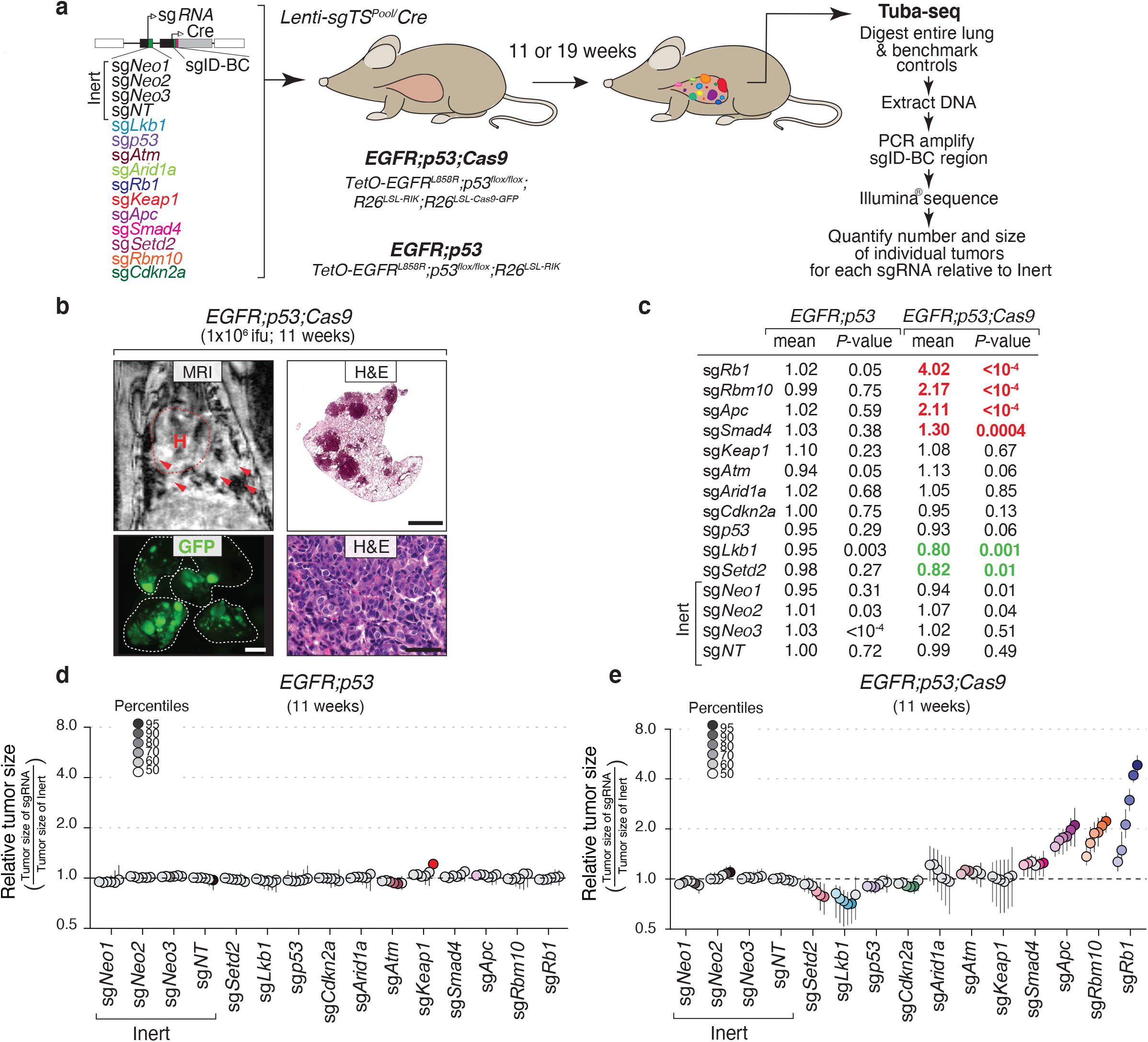
Multiplexed somatic CRISPR–Cas9-mediated genome editing uncovers tumor suppressor gene effects on *EGFR*-driven lung tumors. **a**, Experimental strategy. Tumors were allowed to develop for either 11 weeks in *EGFR;p53;Cas9* and *EGFR;p53* mice or 19 weeks in *EGFR;p53;Cas9* mice after intra-tracheal administration of Lenti-sg*TS*^*Pool*^/*Cre*. Whole lungs were collected for tumor barcode deep sequencing (Tuba-seq) and histology. The number of neoplastic cells in each tumor (tumor size) was calculated from barcode sequencing of bulk tumor-bearing lungs. Barcode read number was normalized to benchmark control cells that have known barcodes and were added at a known number to each sample (Methods). **b**, MRI, H&E, and GFP images showing tumor development in *EGFR;p53;-Cas9* mice 11 weeks after tumor initiation with 1×10^6^ ifu of Lenti-sg*TS*^*Pool*^/*Cre*. H&E image scale bars = 1.2 mm and 100 μm in top and bottom right panels, respectively. Lungs are indicated by the white dashed lines. GFP image scale bar = 2.5 mm. Images are from a representative mouse (*N* = 10). **c**, Relative log-normal (LN) mean size of tumors with each sgRNA in *EGFR;p53* (*N* = 5) and *EGFR;p53;-Cas9* mice (*N* = 10) 11 weeks after tumor initiation (normalized to the tumors with inert sgRNAs). *P*-values were calculated from bootstrap-ping. *P*-values < 0.05 and their corresponding means are highlighted in *EGFR;p53;Cas9* mice for sgRNAs that positively (red) and negatively (green) affect tumor growth when the effects are equal to or differ >10% compared to the size of tumors with inert sgRNAs. **d**, Relative size of tumors at the indicated percentiles of each genotype in *EGFR;p53* mice 11 weeks after tumor initiation with the Lenti-sg*TS*^*Pool*^/*Cre*. These mice lack the *R26*^*LSL-Cas9-GFP*^ allele; therefore, all sgRNAs are functionally inert. 95% confidence intervals are shown. Percentiles calculated from bootstrapping that are significantly different from the tumors with inert sgRNAs are in color. **e**, Relative size of tumors of each genotype in *EGFR;p53;Cas9* mice 11 weeks after tumor initiation with the Lenti-sg*TS*^*Pool*^/*Cre*. The relative size of tumors at the indicated percentiles were calculated from the tumor size distribution of all tumors from ten mice. 95% confidence intervals are shown. Percentiles were calculated from bootstrapping and are in color if significantly different from the tumors with inert sgRNAs.

To assess the function of ten diverse putative tumor suppressor genes, which are frequently altered in human lung adenocarcinoma, we initiated tumors in *EGFR;p53* and *EGFR;p53;Cas9* mice with a pool of barcoded Lenti-sgRNA/*Cre* vectors (Lenti-sg*TS*^*Pool*^/*Cre*; **Fig. 2a**). In addition to Lenti-sgRNA*/Cre* vectors targeting each putative tumor suppressor gene, this pool contains negative control vectors, including four Lenti-sg*Inert/Cre* vectors and a vector with an sgRNA targeting *Trp53* (which is already inactivated by Cre-mediated recombination in *EGFR;p53;Cas9* mice, **Fig. 1a**)^8^. Lenti-sg*TS*^*Pool*^/*Cre*-initiated tumors were first detectable by MRI 4 weeks after tumor initiation in *EGFR;p53;Cas9* mice. At 11 weeks after tumor initiation, when tumors were readily detectable in all mice, tumor-bearing lungs were collected for Tuba-seq analysis and histology (**Fig. 2b** and **Supplemental Fig. 1e**).

### Putative tumor suppressor genes have distinct effects on *EGFR*-driven lung tumor growth

We used Tuba-seq to quantify the number of neoplastic cells in clonal tumors of each genotype in the *EGFR;p53* model and used two summary statistics to describe the tumor size distribution (percentiles within the tumor size distributions and the log-normal mean^34^ (**Methods**; **Figs. 2c-e** and **Supplemental Figs. 2a, b**)). Effects of the sgRNAs on tumor growth were assessed by examining the significance of the differences in the tumor size distribution compared to controls and also the magnitude of the effects. For negative controls, tumors with each sgRNA in *EGFR;p53* mice, which lack the *Cas9* allele, had very similar size distributions (**Fig. 2d**). Furthermore, in *EGFR;p53;Cas9* mice, tumors initiated with each Lenti-sg*Inert/Cre* vector or Lenti-sg*p53/Cre* had very similar tumor size profiles (**Figs. 2c, e**).

Inactivation of *Rb1*, *Apc*, and *Rbm10* most dramatically enhanced the growth of oncogenic *EGFR-*driven *Trp53*-deficient lung tumors (**Figs. 2c, e**). While the importance of *RB1* inactivation in *EGFR*-driven lung adenocarcinomas has begun to be investigated^23,35^, very few studies have addressed the functional importance of the APC pathway on *EGFR*-driven lung adenocarcinomas^34^. Interestingly RBM10 is an RNA binding protein and splicing regulator that is poorly studied in cancer in general and has not previously been implicated as a critical regulator of *EGFR*-driven lung cancer growth^12,36–38^.

Surprisingly, inactivation of either *Lkb1* or *Setd2* – which are strong tumor suppressor genes in analogous oncogenic *Kras*-driven lung tumor models – dramatically reduced tumor growth relative to sg*Inert* tumors^8,39,40^. These effects were consistent across multiple percentiles within the tumor size distribution and as assessed by the LN mean of tumor sizes (**Figs. 2c, e**). Notably, these significant effects observed with sg*Setd2*, sg*Lkb1*, sg*Smad4*, sg*Apc*, sg*Rbm10*, and sg*Rb1* were all recaptured even after we simulated a 50% reduction in cutting efficiency or when we used other strategies for subsampling underscoring the robustness of our findings (**Supplemental Figs. 9b, c** and **Supplemental Figs. 10a, b**). These results suggest that in specific contexts, inactivation of genes presumed to be tumor suppressors can have deleterious effects on cancer growth. Other tumor suppressor genes, *Atm*, *Arid1a*, *Cdkn2a* and *Keap1*, did not significantly alter tumor growth in the context of this experiment (**Figs. 2c, e**).

To assess tumor suppressor gene function at a later time point of tumor growth, we initiated tumors in *EGFR;p53;Cas9* mice with 10-fold less Lenti-sg*TS*^*Pool*^/*Cre* and performed Tuba-seq after 19 weeks of tumor growth. At this time point, the histology of the tumors was similar to that observed after 11 weeks of tumor initiation (**Supplemental Figs. 1c, d**). Tuba-seq analysis confirmed the tumor-suppressive function of *Rbm10*, *Apc,* and *Rb1* (**Supplemental Figs. 2a-c**). Since we used a 10-fold lower viral titer for this experiment, there were proportionally fewer tumors (**Supplemental Fig. 1f**), which limited the resolution of Tuba-seq analysis. Thus, while inactivation of the other genes had no significant effect on tumor growth at this time point, we cannot exclude that these genes may influence tumor growth. Interestingly, despite the decreased statistical power, inactivation of *Cdkn2a* or *Arid1a* had a positive effect on tumor growth at this 19-week time point (but not at the 11-week time point) suggesting a potential role of these tumor suppressor genes during a later phase of tumorigenesis in this model (**Supplemental Figs. 2a, b**).

### Validation of *Apc* and *Rbm10*-mediated tumor suppression

We performed further experiments to confirm the role of two less-well studied tumor suppressors, Apc and Rbm10 on the growth of *EGFR*-driven tumors. We initiated lung tumors in *EGFR;p53* and *EGFR;p53;Cas9* mice with Lenti-sg*Apc*/*Cre*, Lenti-sg*Neo2*/*Cre* (sgInert), and two Lenti-sg*Rbm10/Cre* vectors each with a unique sgRNA targeting *Rbm10* (*N* = 3 *EGFR;p53* mice/group and *N* = 5 *EGFR;p53;Cas9* mice/group; **Fig. 3a**). We used two sgRNAs targeting *Rbm10* to increase the power of our findings, and because the tumor suppressive role of *Rbm10* remains entirely uncharacterized in *EGFR*-driven lung cancer. We found a large variation across mice due to the stochastic nature of tumor progression and the low resolution of the tumor volume measurements determined by MRI and through tumor area calculations (**Supplemental Figs. 3b, c**). Despite this, inactivation of either *Apc* or *Rbm10* in *EGFR;p53;Cas9* mice gave rise to tumors that were significantly larger than control tumors initiated with sg*Neo2* in *EGFR;p53;Cas9* mice. Moreover, by quantifying the size of individual sg*Apc*- or sg*Rbm10*-initiated tumors (based on tumor diameter) in histological sections, we observed that *EGFR;p53;Cas9* tumors were larger than tumors initiated in *EGFR;p53* mice (**Figs. 3b-e**). Lenti-sg*Apc*/Cre-initiated tumors in *EGFR;p53;Cas9* mice had more cancer cells with stabilization and nuclear localization of β-catenin as well as increased expression of Sox9 (consistent with *Apc* inactivation) (**Supplemental Figs. 3d, e**) ^41^. Furthermore, at least 50% of Lenti-sg*Rbm10/Cre*-initiated tumors in *EGFR;p53;Cas9* mice lacked or had lower Rbm10 protein (**Supplemental Fig. 3f**). Tumors with either *Apc* or *Rbm10* inactivation were histologically similar to tumors in *EGFR;p53* mice at this time point and had papillary/acinar or micropapillary structures with a medium/high nuclear grade (**Supplemental Fig. 3a**). These results further confirm the importance of these tumor suppressor pathways in constraining *EGFR*-driven tumor growth *in vivo.* Collectively, our findings underscore the value of coupling Tuba-seq and CRISPR–*Cas9*-mediated somatic genome editing with our virally-induced mouse model to dissect gene function in oncogenic *EGFR*-driven lung cancer.

**Fig. 3.**
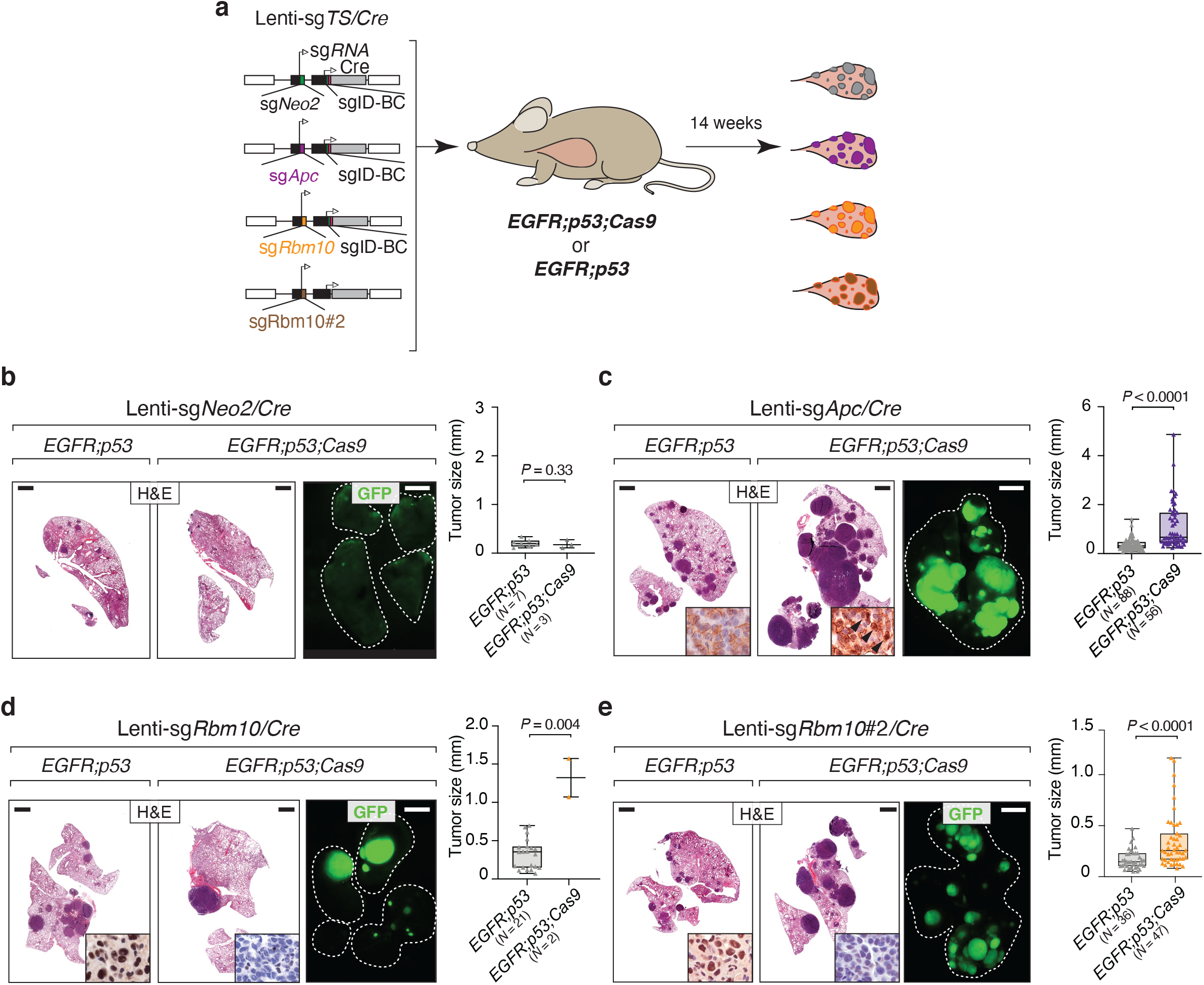
*Apc* and *Rbm10* inactivation enhances the growth of oncogenic *EGFR*-driven lung tumors *in vivo*. **a**, Experimental strategy for tumor suppressor gene validation. Tumors were initiated in *EGFR;p53* and *EGFR;p53;Cas9* mice with Lenti-sg*TS/Cre* vectors carrying sg*Apc*, sg*Rbm10*, sg*Rbm10#2*, or sg*Neo2* as a control (*EGFR;p53 N* = 3 mice/group, *EGFR;p53;Cas9 N* = 5 mice/group, 1×10^5^ ifu/mouse). **b-e**, H&E and GFP images show that *Apc* (**c**) and *Rbm10* (**d, e**) inactivation enhances tumor growth compared to the controls (**b**). The histology of the left and auxiliary lobes was analyzed. Lungs are indicated by the white dashed lines. Scale bars = 1.2 mm and 2.5 mm for histology and GFP images, respectively. Tumor size was calculated by measuring the longest diameter of each tumor. The number of tumors studied are reported on the X-axis. *P*-values were calculated using a one-tailed Mann-Whitney U test. Horizontal lines show the median and whiskers indicate the minimum and the maximum of the data set. The box represents values in the first and the third quartile. The panels on the bottom right of the H&E images show nuclear accumulation of β-catenin (**c**) and absence of Rbm10 protein expression (**d, e**) reflective of *Apc* and *Rbm10* inactivation in tumors in *EGFR;p53;Cas9* mice, respectively.

### Oncogenic drivers shape the fitness landscape of tumor suppression

The extent to which different oncogenic drivers affect the landscape of tumor suppression is almost entirely unknown. We approached this question by comparing the fitness landscape of tumor suppression within the contexts of oncogenic *EGFR-* and *Kras-*driven lung tumors. We repeated an experiment previously performed by our group in which we inactivated the same panel of tumor suppressor genes in *Kras*^*LSL-G12D+*^;*p53*^*flox/flox*^;*R26*^*LSL-Tomato*^;*H11*^*LSL-Cas9*^ (*Kras;p53;Cas9*) mice and used library preparation methods and the analytical pipeline identical to those used for the *EGFR;p53* and *EGFR;p53;Cas9* mice (**Supplemental Fig. 4a**)^29,30,42^. Inactivation of *Lkb1*, *Setd2,* and *Rb1* were particularly strong drivers of oncogenic *Kras*-driven *Trp53*-deficient tumor growth, while inactivation of *Rbm10*, *Apc*, *Cdkn2a* and *Arid1a* also modestly increased tumor growth (**Figs. 4a, b**). These results are largely consistent with our previous Tuba-seq analysis of *Kras;p53;Cas9* mice, as well as other studies on these genes in oncogenic *Kras*-driven lung cancer mouse models (**Supplemental Fig. 1a**)^9,39–41,43–47^

**Fig. 4.**
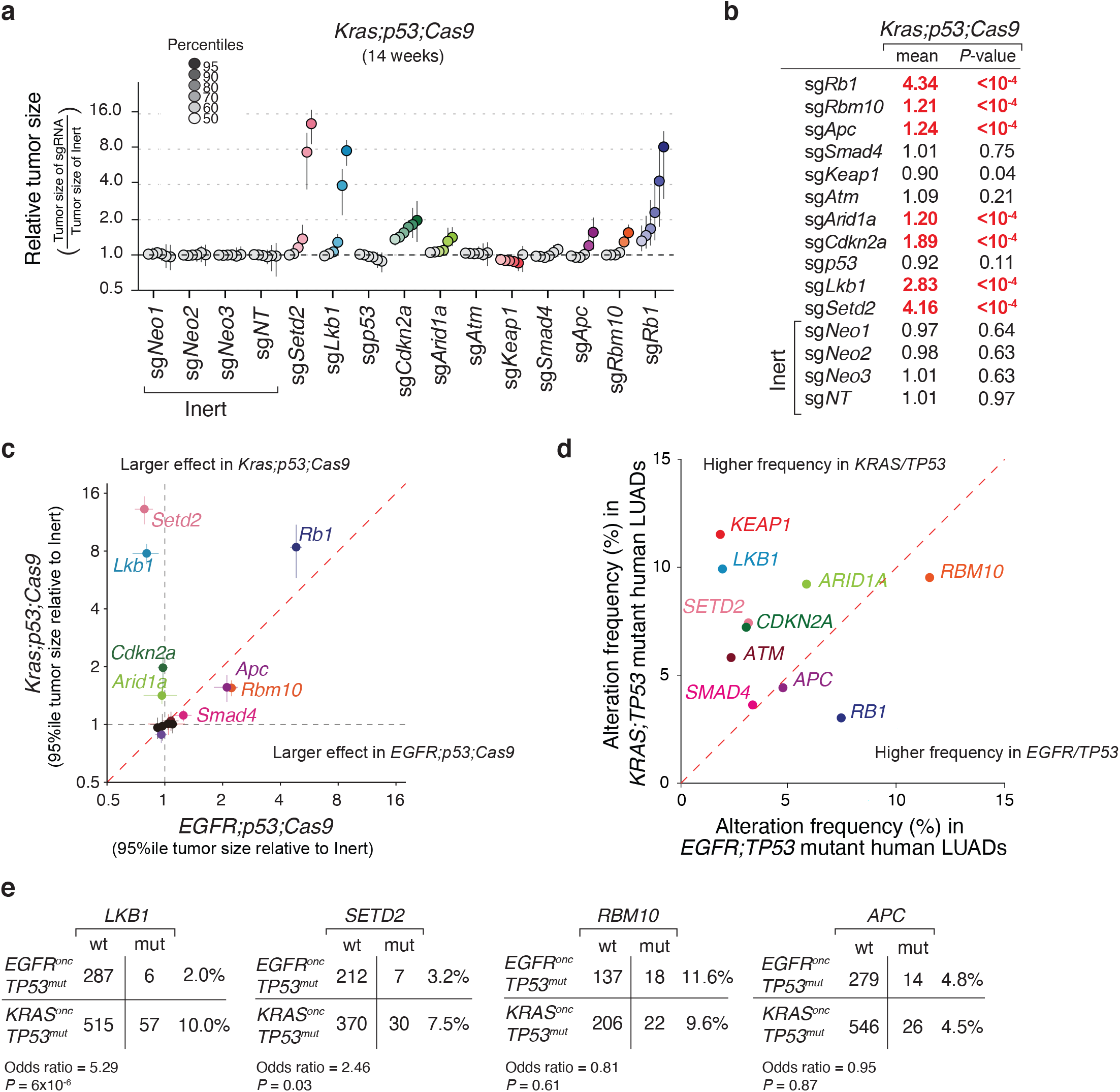
Dramatic differences in the effects of tumor suppressor genes in oncogenic *EGFR-* versus *Kras-*driven lung cancer. **a,** Relative size of tumors of each genotype in *Kras*^*G12D*^;*p53*^*flox/flox*^;*R26*^*LSL-Tomato*^;*H11*^*LSL-Cas9*^ (*Kras;p53;Cas9*) mice 14 weeks after tumor initiation with Lenti-sg*TS*^*Pool*^/*Cre* relative to tumors with inert sgRNAs. The relative size of tumors at the indicated percentiles was calculated from the tumor size distribution of all tumors from six mice. 95% confidence intervals are shown. *P*-values were calculated from bootstrapping. Percentiles that are significantly different from the tumors with inert sgRNAs are in color. **b**, LN mean for tumors with each sgRNA in *Kras;p53;Cas9* mice 14 weeks after tumor initiation (normalized to the inert tumors). *P*-values were calculated by bootstrapping and significant values are highlighted in red when *P* < 0.05 and the effects are >10% compared to the size of tumors with inert sgRNAs. **c**, The relative effect of inactivating each tumor suppressor gene on tumor sizes (comparison to tumors with inert sgRNAs) at the indicated percentile in the *EGFR;p53;Cas9* (*N* = 10) and *Kras;p53;Cas9* (*N* = 6) models. *Lkb1* and *Setd2* inactivation greatly increased tumor size only in the *Kras;p53;Cas9* model. Error bars indicate the standard deviation. **d**, The frequency of tumor suppressor gene alterations that co-occur with *EGFR* or *KRAS* and *TP53* mutations in human lung adenocarcinomas (LUADs, data from AACR Project GENIE). **e**, *LKB1* and *SETD2* alterations co-occur significantly more frequently in *KRAS*^*onc*^;*TP53*^*mut*^ than in *EGFR*^*onc*^;*TP53*^*mut*^ human lung adenocarcinomas. The frequency of *RBM10* and *APC* alterations were not significantly different between in *KRAS*^*onc*^;*TP53*^*mut*^ and *EGFR*^*onc*^;*TP53*^*mut*^ tumors. The odds ratios represent the strength of the observed frequencies of tumor suppressor gene alterations that co-occur in *KRAS*^*onc*^;*TP53*^*mut*^ cases compared to *EGFR*^*onc*^;*TP53*^*mut*^ cases. *P*-values were calculated using a Fisher’s exact test.

Inactivation of several of the tumor suppressor genes (e.g. *Rb1*) had similar effects on *EGFR-* and *Kras-*driven tumors suggesting that these putative tumor suppressor genes limit lung adenocarcinoma growth regardless of the oncogenic context in these mouse models (**Fig.4c** and **Supplemental Fig. 4b**). However, inactivation of either *Lkb1* or *Setd2* greatly increased the growth of oncogenic *Kras*-driven lung tumors but decreased the growth of oncogenic *EGFR*-driven lung tumors (**Figs. 2c, e**, **Figs. 4a-c** and **Supplemental Fig. 4b**). Thus, the consequences of tumor suppressor gene inactivation in specific contexts are not limited to the magnitude of tumor-suppressive effects but can also be manifested as opposite effects (known as the sign epistasis) even when the driving oncogenic alterations (in this case *EGFR* and *KRAS*) are traditionally thought to be within a linear pathway.

### Profound epistasis between tumor suppressor genes and oncogenic drivers drives mutational patterns in human lung adenocarcinoma

To compare our functional data from our *in vivo* models with the spectrum of tumor suppressor gene mutations found in human lung adenocarcinomas, we queried data from the AACR Project GENIE database^48^. We calculated the frequency of tumor suppressor gene mutations that occur co-incident with oncogenic *EGFR* (L858R, Exon 19 deletions, L861Q, G719X) or oncogenic *KRAS* (at codons 12, 13 or 61) mutations and *TP53* mutations. This analysis revealed different frequencies of mutations in several tumor suppressor genes in *EGFR/TP53* and *KRAS/TP53* mutant human lung tumors. *RB1*, *RBM10*, and *APC* are frequently altered tumor suppressor genes in *EGFR/TP53* mutant lung adenocarcinomas. Interestingly, *RB1* mutations are more frequent in *EGFR/TP53* tumors compared to *KRAS/TP53* tumors (7.5% versus 3.1%). However, *Rb1* inactivation was a major driver of tumor growth in both *EGFR* and *Kras* mutant tumors in mice (**Figs. 2c, e** and **Figs. 4a-c**). This apparent discrepancy between mouse and human may be related to the higher frequency of alterations in *CDKN2A* in human *KRAS/TP53* mutant tumors (7.3%, **Fig. 4d**) that would disrupt the same cell cycle regulation pathway as *RB1* inactivation. *LKB1/STK11* and *SETD2,* are among the most frequently mutated tumor suppressor genes in *KRAS/TP53* mutant lung adenocarcinomas (**Fig. 4d**)^49,50^. Further supporting a difference in the function of *LKB1* and *SETD2* in *EGFR/TP53* versus *KRAS/TP53* mutant lung adenocarcinomas, mutations in these genes occurred at significantly higher frequencies in *KRAS/TP53* mutant tumors compared to *EGFR/TP53* tumors (**Fig. 4e**). This asymmetry in the mutation frequency of *LKB1* or *SETD2* within oncogenic *EGFR-* or *KRAS*-driven lung tumors is also significant when we extend our analysis to include all tumors regardless of *TP53* mutation status (**Supplemental Fig. 4c**). Collectively, the mouse and human data indicate that the mutational patterns can reflect the biological consequences of profound epistasis between tumor suppressor genes and oncogenic drivers. This highlights the power of our approach in uncovering the functional relevance of genomic combinations on tumorigenesis.

### *Keap1* inactivation limits the response of tumors to osimertinib

Genetically engineered mouse models have provided insight into the biology of *EGFR*-driven lung tumors and proven valuable in studying mechanisms of resistance to EGFR TKIs, especially on-target ^51^ mechanisms of resistance^27,52–54^. The TKI osimertinib was recently approved for the first-line treatment of *EGFR*-driven lung adenocarcinomas. However, pathways involved in modulating the depth of response and mechanisms of resistance to osimertinib are still under investigation^55,56^. To investigate how tumor suppressor genes influence the therapeutic response of lung tumors to EGFR inhibition, we treated *EGFR;p53;Cas9* mice with Lenti-sg*TS^Pool^/Cre*-initiated tumors with osimertinib for two weeks starting at 9 weeks after tumor initiation (**Fig. 5a**). Osimertinib treatment greatly reduced the overall tumor burden relative to vehicle-treated *EGFR;p53;Cas9* mice (**Supplemental Figs. 5a-d**). Residual neoplastic cells were sparse, as determined by staining for EGFR^L858R^ and those cells were not proliferating (**Supplemental Figs. 5e-f**). The overall tumor response was similar when the 2-week treatment was started 17 weeks after tumor initiation (**Supplemental Figs. 5g-j**).

**Fig. 5.**
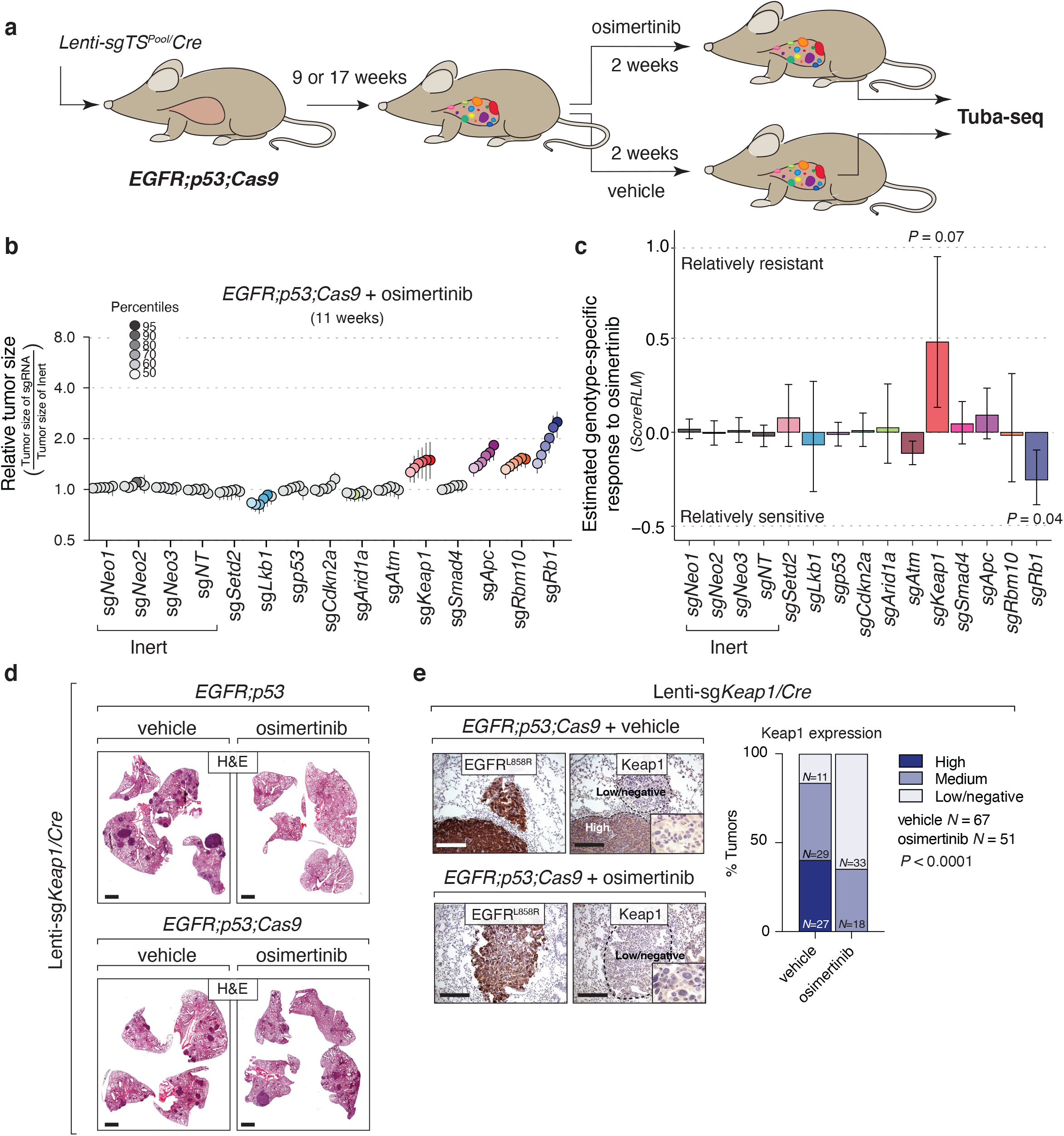
Identification of genotype-specific responses to osimertinib treatment. **a,** Experimental strategy. Tumors were initiated with Lenti-sg*TS*^*Pool*^/*Cre* in *EGFR;p53;Cas9* mice for nine or 17 weeks and were treated for two weeks with either vehicle or osimertinib (25 mg/kg, 5 days/week). **b**, Relative size of the tumors at the indicated percentiles for each genotype in *EGFR;p53;Cas9* mice after treatment with osimertinib. 95% confidence intervals are shown. *P*-values were calculated from bootstrapping. Percentiles that are significantly different from the tumors with inert sgRNAs are in color. **c**, Estimate of the genotype-specific treatment response (*ScoreRLM*) calculated by comparing the LN mean of tumors treated with osimertinib to the LN mean of vehicle-treated tumors in *EGFR; p53;Cas9* mice 11 weeks after tumor initiation (Methods). Error bars indicate the standard deviation. *P*-values were calculated from bootstrapping. **d**, H&E staining of tumor bearing-lungs in *EGFR;p53* and *EGFR;p53;Cas9* mice with Lenti-sg*Keap1/Cre* initiated tumors (*N* = 8 mice/group). Scale bars = 1.2 mm. **e**, Immunostaining of Lenti-sg*Keap1/Cre* initiated tumors in *EGFR;p53;Cas9* mice for EGFR^L858R^ and Keap1 after vehicle or osimertinib treatment (*N* = 4 mice/treated-group). Tumors are still detectable after two weeks of treatment with osimertinib and mostly express medium or low/negative levels of Keap1 compared to tumors treated with vehicle in *EGFR;p53;Cas9* mice (bar graph). The dashed lines indicate areas of tumors and the level of Keap1 is indicated with a label (high or low/negative). Scale bars = 200 μm. *P*-values were calculated using a Chi-squared test.

To quantify the impact of inactivating each tumor suppressor gene on the response to osimertinib *in vivo*, we performed Tuba-seq on the lungs from osimertinib-treated *EGFR;p53;Cas9* mice 11 and 19 weeks after tumor initiation and compared the results to Tuba-seq results from vehicle-treated controls (**Fig. 5b** and **Supplemental Figs. 6a, b**). Consistent with the imaging data and histological analysis, osimertinib treatment greatly reduced tumor burden as assessed by Tuba-seq (**Compare Figs 2c, e with Fig. 5b and Supplemental Fig. 6a**; **Methods**).

After two weeks of osimertinib treatment, inactivation of *Apc*, *Rb1* or *Rbm10* was still associated with larger tumors while the size distribution of tumors with inactivation of *Cdkn2a*, *Arid1a*, or *Atm* remained similar to that of tumors with inert sgRNAs (compare **Fig. 5b** and **Fig. 2e, Supplemental Fig. 6a** and **Fig. 2c**, **Supplemental Fig. 6b** and **Supplemental Fig. 2a**). The striking exception was tumors with sg*Keap1*. In vehicle-treated mice, the size distribution of sg*Keap1* tumors was almost identical to that of tumors with inert sgRNAs, however in osimertinib-treated mice sg*Keap1* tumors were significantly larger than the tumors with inert sgRNAs. This suggests that inactivation of *Keap1* limits responses to osimertinib (**Fig. 5b**). Osimertinib resistance conferred by *Keap1* inactivation was also observed at 19 weeks after tumor initiation (**Supplemental Figs. 6a, b**).

We applied an analytical approach that we previously developed and validated to quantify the genotype-specific responses (**Fig. 5c** and **Supplemental Figs. 6c-g**; **Methods**)^57^. By comparing the LN mean of the observed tumor size distributions in osimertinib-treated mice with the expected tumor size distribution based on the overall drug effects, we can estimate genotype specific drug responses (*ScoreRLM*; **Fig. 5c** and **Supplemental Fig. 6c**). At 11 weeks after tumor initiation, following 2 weeks of osimertinib treatment sg*Rb1* tumors were 25% smaller than expected (*P* = 0.04). Conversely, tumors with *Keap1* inactivation were 48% larger than expected (*P* = 0.07; **Fig. 5c**). The effect of *Keap1* inactivation was even greater at 19 weeks after tumor initiation, where tumors were 274% larger than expected (*P* = 0.13, **Supplemental Fig. 6c**). Given the magnitude of the *ScoreRLM* for sg*Keap1* at both time points (*ScoreRLM* = 0.57 and 1.90 after two weeks of treatment at 11 and 19 weeks after tumor initiation, respectively), we combined the two independent *P*-values and confirmed that *Keap1* inactivation significantly reduced the therapeutic response to osimertinib (Fisher’s method, *P*-value = 0.05). Other statistical measures of genotype-specific responses, including relative tumor number (*ScoreRTN*) and relative geometric mean (*ScoreRGM*) did not significantly differ between the treated and untreated groups (**Supplemental Fig. 6d**; **Methods**). Our analytical methods allow us to uncover when effects are more pronounced on larger tumors (*ScoreRTN* and *ScoreRGM* have much lower sensitivity when the effects are greater on larger tumors; **Supplemental Figs. 6e-g**; **Methods**). Thus, our data are consistent with the resistance conferred by *Keap1* inactivation being more pronounced in larger tumors.

### *Keap1*-deficient tumors have reduced sensitivity to osimertinib which correlates with clinical outcomes

To further investigate these findings, we initiated tumors with Lenti-sg*Keap1/Cre* in *EGFR;p53* and *EGFR;p53;Cas9* mice followed by treatment with osimertinib or vehicle (**Figs. 5d, e**). Osimertinib-treatment reduced the size and number of tumors in *EGFR;p53* mice compared to vehicle-treated *EGFR;p53* mice (**Fig 5d** and **Supplemental Figs. 7a, b**). Conversely, osimertinib treatment of Lenti-sg*Keap1/Cre*-initiated tumors in *EGFR;p53;Cas9* mice did not decrease tumor size or number compared to vehicle-treated *EGFR;p53;Cas9* mice (**Figs. 5d, e** and **Supplemental Figs. 7a, b**). Consistent with the inefficiency of CRISPR–Cas9-mediated genome editing in somatic cells, some tumors initiated with Lenti-sg*Keap1/Cre* in *EGFR;p53;Cas9* mice retained expression of Keap1 protein (**Fig. 5e** top panels). However, the tumors that remained in osimertinib-treated *EGFR;p53;Cas9* mice all had medium to low/negative expression of Keap1 (**Fig. 5e**). Together, these data indicate that while *Keap1* inactivation is not positively selected for during oncogenic *EGFR*-driven tumor growth, osimertinib treatment selects for cancer cells expressing low/negative levels of *Keap1*, thus reducing the therapeutic response to the drug.

To correlate our findings with clinical data, we analyzed the effects of KEAP1 pathway alterations on patient outcomes to EGFR inhibition in *EGFR/TP53* mutant lung adenocarcinomas. Oncogenic *EGFR-*driven tumors with KEAP1 pathway inactivation have been suggested to be less responsive to TKIs, and we confirmed this association in *EGFR/TP53* tumors (**Figs. 6a, b**)^58^. Mutations in the *KEAP1/NFE2L2/CUL3* pathway were associated with a significantly shorter time on EGFR TKI therapy compared with matched patients with *KEAP1/NFE2L2/CUL3* wildtype tumors (5.8 versus 14.3 months, *P* = 0.01; Log-rank test, **Fig. 6a**; **Supplemental Table 1**). This remained significant even after adjustment for potential confounders such as age, sex, race and smoking status (**Supplemental Table 2**). Among several other tumor suppressor genotypes, KEAP1 pathway alterations were the most significant driver of limited sensitivity after correction for multiple hypothesis testing (**Fig. 6b**).

**Fig. 6.**
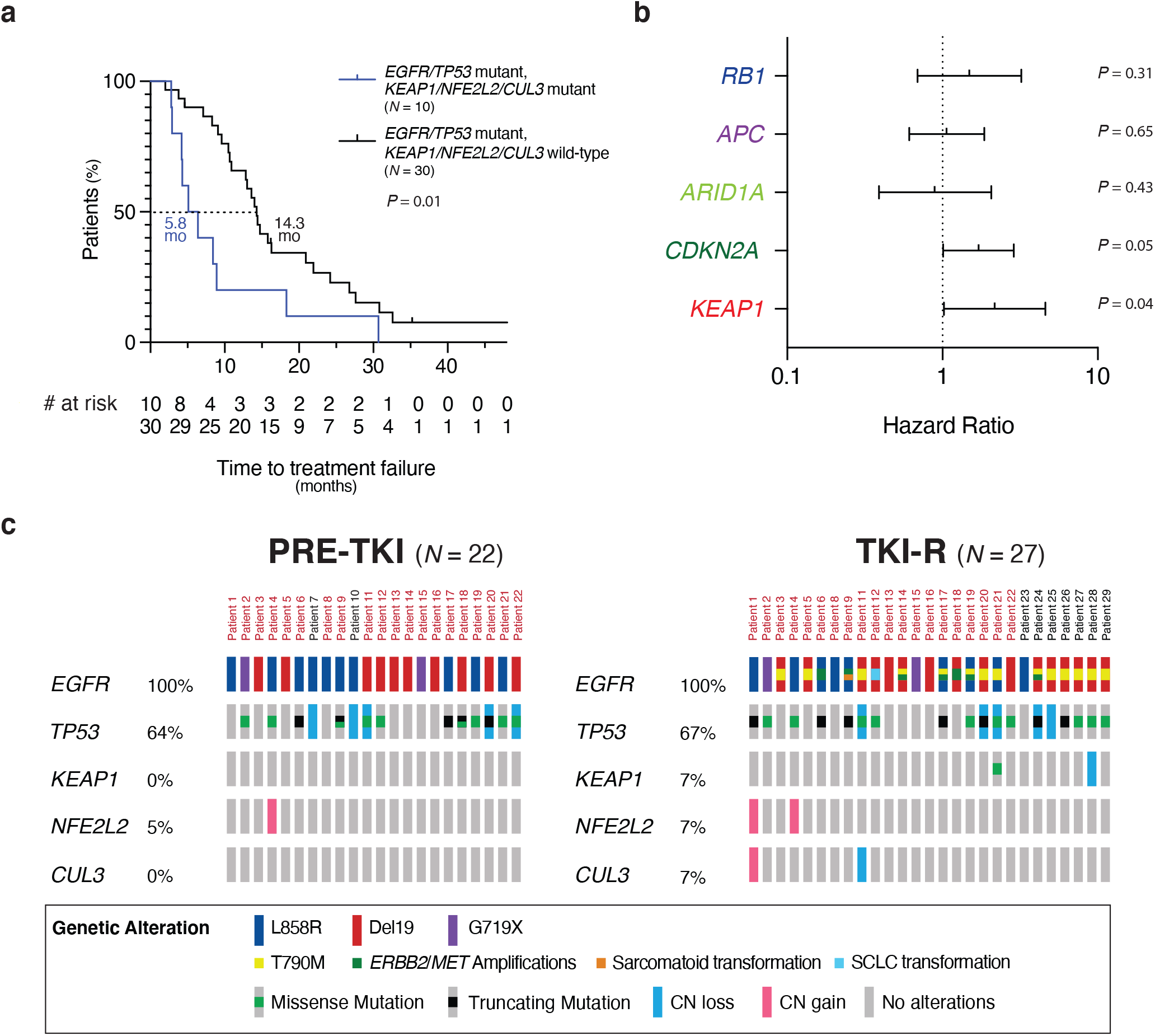
*KEAP1* inactivation correlates with reduced therapeutic response to TKIs in human *EGFR*-driven lung adenocarcinomas. **a**, Kaplan-Meier curve showing time to TKI treatment failure for patients with *EGFR/TP53* lung adenocarcinomas that do or not have mutations in the *KEAP1/NFE2L2/CUL3* axis. Time to treatment failure is the time from the initiation of TKI treatment to the date of discontinuation of TKI due to progression, toxicity or death. The *P*-value was calcuated using the Log rank test. **b**, Forest plot of the time on treatment hazard ratios for tumors with *KEAP1*, *RB1*, *APC*, *ARIDA* or *CDKN2A* alterations in *EGFR/TP53* mutant lung adenocarcinomas. Hazard ratios were calculated from multivariate regression analysis, 95% confidence intervals are shown. **c**, *KEAP1, NFE2L2*, and *CUL3* alteration frequency in oncogenic *EGFR* and *TP53* mutant lung tumors in the Yale Lung Rebiopsy Program dataset (Methods). OncoPrint of oncogenic *EGFR* samples prior to first-line TKI (erlotinib, gefitinib, afatinib) treatment (PRE-TKI, left panel) (*N* = 22), and at acquired resistance to TKIs (TKI-R, right panel) (*N* = 27). In each row, the percentage and the type of alterations in each gene are indicated for each patient. For the TKI-R samples, known resistance mechanisms such as acquisition of T790M in *EGFR*, are reported. Cases with paired PRE-TKI and TKI-R samples are indicated by the patient number highlighted in red.

We also analyzed a dataset of oncogenic *EGFR* lung adenocarcinoma patient samples collected through the Yale Lung Rebiopsy Program (YLR) prior to first-line TKI treatment (erlotinib, gefitinib, or afatinib) and after the development of resistance to TKIs (**Fig. 6c**; **Supplemental Table 3**). Among 18 patients with *EGFR* and *TP53* mutant tumors, two had *KEAP1* alterations at relapse (TKI-R samples). One case had an acquired missense mutation (Y206N) that lies in a domain of KEAP1 involved in forming the complex with Cullin3 to mediate ubiquitination and degradation of NRF2 (encoded by *NFE2L2*)^59^. The other tumor, which was analyzed only at resistance had heterozygous loss of *KEAP1* (**Fig. 6c**). Moreover, we also observed two cases of *NFE2L2* copy number gain (one detected prior to treatment and maintained at resistance, the other detected only at resistance), and one case of *CUL3* heterozygous loss at resistance. Thus, our *in vivo* functional results are consistent with human data and support a role for inactivation of the KEAP1 pathway in reducing EGFR TKI sensitivity.

## Discussion

The genomic fitness landscape of lung adenocarcinoma is combinatorially complex, with a large number of genomic alterations in oncogenic drivers and tumor suppressor genes occurring in various combinations^2,3^. While critical for precision medicine approaches, dissecting the functional contribution of these co-occurring genetic alterations to tumor growth and response to therapy is extremely challenging. Using somatic CRISPR–Cas9-mediated gene editing and Tuba-seq^8,9^, we evaluated the fitness effects of inactivation of ten tumor suppressor genes commonly altered in lung adenocarcinoma in the contexts of oncogenic *EGFR*- and *Kras*-driven tumors. We uncovered distinct roles for specific tumor suppressor genes on tumor growth in the different oncogenic contexts and on sensitivity to the TKI osimertinib *in vivo*.

*EGFR* and *KRAS* are the most frequently mutated oncogenic drivers in lung adenocarcinoma, collectively occurring in ~40-50% of all patients^2^. Oncogenic alterations in these two genes are mutually exclusive, consistent with them being in the same canonical receptor tyrosine kinase (EGFR-RAS-MAPK) signaling pathway^60,61^. For this reason, one might anticipate that inactivating tumor suppressor pathways in *EGFR*-driven and *KRAS*-driven lung cancers would have similar biological consequences. It is tempting to make a simplifying assumption that tumor suppressor genes have a constant marginal effect independent of other genomic alterations, or in other words, that the fitness landscape of tumor suppression is smooth and that epistatic interactions are infrequent and/or small in magnitude. Here, we tested this assumption by comparing the fitness landscape of tumor suppression in *EGFR*-driven *Trp53*-deficient and in *Kras*-driven *Trp53*-deficient lung adenocarcinoma and found pervasive epistasis between tumor suppressor genes and oncogenes. Inactivation of *Rb1*, *Rbm10* and *Apc* had a similar effect on *EGFR* and *KRAS*-driven lung tumors (**Figs. 2c, e**, **Figs. 4a-c** and **Supplemental Figs. 4b**). However, while *Lkb1* and *Setd2* were amongst the most potent tumor suppressor genes in *Kras/p53* tumors, sg*Lkb1* and sg*Setd2* led to reduced tumor growth of *EGFR/p53* tumors (**Figs. 2c, e** and **Figs. 4a, b**). This observation is consistent with human data, where alterations in either *LKB1* or *SETD2*, are much less common in lung tumors with oncogenic *EGFR* than in tumors with oncogenic *KRAS* (**Figs. 4d, e** and **Supplemental Fig. 4c**). Thus, we reveal a highly context-dependent fitness landscape of tumor suppression that depends on the nature of the oncogenic driver. Investigation of the mechanisms that underlie the sign epistasis of *LKB1* and *SETD2* may uncover new biological insights and vulnerabilities of *EGFR* mutant tumors. More broadly, sign epistasis within seemingly similar cancer contexts, could help identify genetic interactions for further functional investigation and should be considered when interpreting cancer genomic data (**Figs. 4d, e** and **Supplemental Fig. 4c**).

One major advantage of autochthonous genetically engineered mouse models of human cancer is that they can be used to study the impact of inactivation of putative tumor suppressor genes on therapy responses *in vivo*^10,57,62^. We found that inactivation of *Keap1* decreases sensitivity to osimertinib *in vivo* (**Fig. 5**). This finding was supported in a clinical cohort of *EGFR/TP53* mutated lung adenocarcinomas, in which patients with tumors harboring mutations in *KEAP1/NFE2L2/CUL*3 had a significantly shorter time to treatment discontinuation with EGFR TKIs compared with matched controls (**Figs. 6a, b** and **Supplemental Figs. 8a-d**)^17,58^. These results are consistent with this tumor suppressor gene pathway limiting the response to therapy and explains the presence of alterations in *KEAP1* pathway genes in TKI-resistant human tumors. A reduced response to TKI therapy mediated by *KEAP1* inactivation may be associated with the accumulation and transactivation of oxidative stress-related genes by NRF2^63–65^. Indeed, we found that Keap1-deficient tumors had increased nuclear Nrf2 and Nqo1 expression relative to Keap1-proficient tumors, suggesting enhanced Nrf2 transcriptional activity (**Supplemental Fig. 7c**)^63–65^. Alternatively, KEAP1/NRF2-dependent metabolic reprogramming could be involved in mediating drug resistance in lung cancer^66–68^. However, direct genetic alterations in this pathway occur in less than 10% of TKI-resistant *EGFR* mutant lung adenocarcinomas (**Fig. 6c**)^17^. It is possible that non-genomic alterations that increase NRF2-dependent gene expression or other mechanisms that decrease oxidative stress also occur in TKI-resistant tumors^69–72^. Collectively, these results raise the possibility that targeting NRF2 may reduce or delay the onset of resistance in *EGFR*-driven lung adenocarcinomas. More broadly, our study demonstrates that our approach can be used to identify clinically relevant pathways that modulate response to therapy *in vivo*. By uncovering the driving forces of the heterogeneity of responses to therapy observed in patients, these types of studies could help define high-risk versus low-risk patient populations and guide therapeutic interventions.

This study provides insight into the complex interplay between tumor suppressor genes and other co-occurring mutations in *EGFR*-driven lung adenocarcinoma tumorigenesis and thus has significant clinical implications. By evaluating interactions between co-occurring alterations in these models, we have avoided confounding factors pervasive in human genomic data (*i.e.,* tumor mutation load, mutation frequency, passenger mutations) and environmental factors such as smoking, a condition that is more often, but not exclusively associated with *KRAS*-driven tumors. Our data provide clear quantitative data on mutual exclusivity and synergistic biological effects of genetic alterations. Notably, other oncogenic drivers (e.g. an *ALK* rearrangement) also have a unique spectrum of co-occurring tumor suppressor gene alterations further suggesting wide-spread interactions between tumor suppressor gene pathways and oncogenic drivers (**Supplemental Fig. 11**). Future *in vivo* Tuba-seq studies should investigate tumor models driven by other oncogenes to uncover a broader understanding of the genetic interactions between diverse oncogenes and large panels of tumor suppressor genes. Precise mapping of the fitness consequences of combinations of genetic alterations during tumor evolution will help uncover the biological and clinical relevance of specific alterations during carcinogenesis and identify pathways that can be exploited as therapeutic targets to prevent or overcome resistance to TKIs.

## Acknowledgements

G.F was supported by an American Italian Cancer Foundation (AICF) postdoctoral fellowship. C.L. is the Connie and Bob Lurie Fellow of the Damon Runyon Cancer Research Foundation (DRG-2331). H.C. was supported by a Tobacco-Related Disease Research Program Postdoctoral Fellowship (28FT-0019). D.A. was supported by NIH F31 CA203488. K.H. was supported by NIH F32 CA210516. This work was supported by NIH R01 CA231253 (to M.M.W and D.P), NIH R01 CA234349 (to M.M.W and D.P.), NIH R01 CA120247 (to K.P.) and NIH P50 CA196530 (to K.P.). The authors would like to acknowledge the American Association for Cancer Research and its financial and material support in the development of the AACR Project GENIE registry, as well as members of the consortium for their commitment to data sharing. Interpretations are the responsibility of study authors. We thank members of the Politi, Winslow and Petrov Laboratories for helpful feedback and support in particular, Jacqueline Starrett and Emily Forrest for editing, Fernando de Miguel for creating resources to help with the WES analysis. We also thank Henning Stehr for providing the sequencing data from the STAMP cohort and the Bosenberg Laboratory for sharing their Leica M205 FA stereomicroscope.

## Author contributions

G.F. designed and performed *in vivo* experiments, analyzed the data and wrote the manuscript. C.L. performed sequencing analysis, developed and performed statistical analyses, and wrote the manuscript. H.C. designed sgRNAs, generated Lenti-sgRNA*/Cre* vectors, tested sgRNA cutting efficiency, produced lentivirus, and wrote the manuscript. J.A.H. evaluated and analyzed the Stanford Solid Tumor Actionable Mutation Panel (STAMP) cohort of lung cancer patients at Stanford University School of Medicine and wrote the manuscript. W.L. performed *in vivo* experiments, D.A. and K.H. developed the models used for the study. J.C. generated the pipeline for the WES data of the Yale Rebiopsy program (YLR) cohort at the Yale Cancer Center. A.W. coordinates the YLR program. L.A. prepared the libraries for Tuba-seq. D.M. developed the pipeline to analyze the GENIE data. N.R. and S.L. maintained the Yale mouse colony and imaged mice included in this study. R.H. performed the histological analysis of all the mouse lung tumor samples. S.G. conceptualized and leads the Yale Rebiopsy Program at the Yale Cancer Center with K.P.. M.D. conceptualized and analyzed the STAMP cases. H.W. supervised the analysis of the STAMP cohort. D.A.P. conceptualized, supervised the project and wrote the manuscript. M.M.W. and K.P. designed the experiments, conceptualized, supervised the project and wrote the manuscript. All the authors have revised and approved the submitted version.

## Methods

### Mice and tumor initiation

*TetO-EGFR*^*L858R*^; *p53*^*flox/flox*^; *Rosa26*^*CAGs-LSL-rtTA3-IRES-mKate*^, *Rosa26*^*CAGs-LSL-Cas9-GFP*^, *Kras*^*LSL-G12D*^, *Rosa26*^*LSL-tdTomato*^, and *H11*^*LSL-Cas9*^ mice have been described^4,25,27–30,42,73,74^. *EGFR*;*p53* and *EGFR*;*p53;Cas9* were on a mixed BL6/129/FVB background and *Kras*;*p53;Cas9* mice were on a mixed BL6/129 background. Approximately equal numbers of males and females were used for each experiment and the number of mice used for each experiment is listed in each figure legend. Lung tumors were initiated by intratracheal administration of Lentiviral-*Cre* vectors as previously described^26^. Tumor burden was assessed by magnetic resonance imaging, fluorescence microscopy, lung weight, and histology, as indicated. Doxycycline was administered by feeding mice with doxycycline-impregnated food pellets (625 ppm; Harlan-Teklad). Osimertinib (from AstraZeneca, Cambridge, UK) was resuspended in 0.5% (w/v) methylcellulose (vehicle) and was administered orally (*per os*, 25 mg/kg 5 days a week). All animals were kept in pathogen-free housing under guidelines approved by either the Yale University Institutional Animal Care or the Stanford University Institutional Animal Care and Use Committee guidelines.

### Production, purification, and titering of lentivirus

The barcoded vectors in the Lenti-sg*TS*^*Pool*^/*Cre* have been previously described (**Supplemental Table 4**)^8^. The second Lenti-sg*Rbm10/Cre* vector used in the validation experiments was generated by site-directed mutagenesis (**Supplemental Table 5**). Briefly, Lentiviral-U6-sgRNA*/Cre* vectors contain an 8-nucleotide defined sequence (sgID) that identifies the sgRNA followed by a 15-nucleotide random barcode (BC) to uniquely tag each tumor^8^. To avoid barcode-sgRNA uncoupling driven by lentiviral template switching during reverse transcription of the pseudo-diploid viral genome, each barcoded Lenti-sgRNA*/Cre* vector was generated separately^75,76^. We cultured HEK293T cells in Dulbecco’s Modified Eagle Medium with 10% Fetal Bovine Serum and transfected them with individual barcoded Lenti-sgRNA*/Cre* plasmids (*sgLkb1, sgp53, sgApc, sgAtm, sgArid1a, sgCdkn2a, sgKeap1, sgNeo1, sgNeo2, sgNeo3, sgNT1, sgRb1, sgRbm10, sgRbm10#2* unbarcoded*, sgSetd2,* or *sgSmad4*) along with pCMV-VSV-G (Addgene #8454) envelope plasmid and pCMV-dR8.2 dvpr (Addgene #8455) packaging plasmid using polyethylenimine. We treated the cells with 20 mM sodium butyrate 8 hours after transfection, changed the culture medium 24 hours after transfection, and collected supernatants 36 and 48 hours after transfection. Subsequently, we removed the cell debris with a 0.45 μm syringe filter unit (Millipore SLHP033RB), concentrated each lentiviral vector by ultracentrifugation (25,000 g for 1.5 hours at 4°C), resuspended the virus in PBS, and stored the virus at −80°C. To determine the titer of each vector, we transduced *Rosa26*^*LSL-YFP*^ mouse embryonic fibroblasts (a gift from Dr. Alejandro Sweet-Cordero/UCSF), determined the percentage of YFP-positive cells by flow cytometry, and normalized the titer to a lentiviral preparation of known titer. Lentiviral vectors were thawed and pooled immediately prior to delivery to mice. All these plasmids are available at https://www.addgene.org/Monte_Winslow/.

### Lentiviral titers and time of analysis

Anticipated growth rates were determined by monitoring tumor development through magnetic resonance imaging in pilot experiments and the analysis time points were selected to ensure that tumors were detectable by MRI such that their response to treatment could be evaluated. Viral titers used in the experiments were chosen to balance the total tumor burden across mice at the time of analysis after tumor initiation. For analysis of tumor growth 11 weeks after tumor initiation, the Lenti-sg*TS*^*Pool*^/*Cre* titer administered to *EGFR*;*p53* mice was 2×10^6^ infectious units (ifu)/mouse, while for *EGFR*;*p53;Cas9* mice we used 1×10^6^ ifu/mouse. We reasoned that using a higher viral titer in the control *EGFR;p53* mice would increase our confidence that any differences observed between *EGFR;p53* and *EGFR;p53;Cas9* mice were due to inactivation of tumor suppressor genes in the latter model. For the 19-week time point in *EGFR*;*p53;Cas9* mice we initiated tumors with 1×10^5^ ifu/mouse. Two weeks before collection, mice were treated with either vehicle or osimertinib. For the validation experiments in which we used a single vector to initiate tumors (Lenti-sg*Apc/Cre*, Lenti-sg*Rbm10/Cre,* Lenti-sg*Rbm10*#2*/Cre* or Lenti-sg*Neo2/Cre* (sg*Inert*)) we used 1×10^5^ ifu/mouse and analyzed the mice after 14 weeks of tumor growth (**Supplemental Table 5**). For the validation with Lenti-sg*Keap1/Cre* virus (1×10^5^ ifu/mouse), both *EGFR;p53* and *EGFR;p53;Cas9* mice were treated 15 weeks after tumor initiation and lungs were collected after two weeks of treatment with either vehicle or osimertinib. *Kras*;*p53;Cas9* were analyzed 14 weeks after tumor initiation with 2.2×10^4^ ifu/mouse.

### Magnetic resonance imaging

All procedures were performed in accordance with protocols approved by the Yale University IACUC and in agreement with the NIH Guide for the Care and Use of Laboratory Animals. Respiratory gated, gradient-echo MR images of mice were collected with a 4T (31-cm bore) small-animal Bruker horizontal-bore spectrometer (Bruker AVANCE). All data were collected as previously described^52^. Tumor volume was quantified by calculating the area of visible lung opacities present in each image sequence per mouse using BioImage Suite 3.01^77^.

### Isolation of genomic DNA from mouse lungs and preparation of sgID-BC libraries

Genomic DNA was isolated from bulk tumor-bearing lung tissue from each mouse as previously described^8^. Briefly, three benchmark control cell lines (~5×10^5^ cells per cell line) carrying unique sgID-BCs, were added (“spiked-in”) to each sample prior to lysis to enable the calculation of the absolute number of neoplastic cells in each tumor from the number of sgID-BC reads. Following homogenization and overnight protease K digestion, genomic DNA was extracted from the lung lysates using standard phenol-chloroform and ethanol precipitation.

sgID-BC sequencing libraries were prepared by PCR amplifying the sgID-BC region from total genomic DNA. To enable the identification and subsequent computational elimination of index hopped reads after high-throughput sequencing, the sgID-BC region of the integrated Lenti-sg*RNA-BC/Cre* vectors was PCR amplified using unique dual indexing primer pairs. To increase the sequence diversity at each position and reduce the amount of PhiX required to achieve high sequencing quality, we added 6-9 Ns before the sequence-specific primers^57^. PhiX is control DNA (with a diverse sequence) that is added to Illumina sequencing samples when the diversity of the product to be sequenced is low (often the case with amplicon sequencing like what we are doing to analyze barcodes). Since we are performing amplicon sequencing with the possibility of less than random sequence diversity at each position, we add 5%-15% PhiX to ensure good sequencing quality. We used a single-step PCR amplification of sgID-BC regions, which we have found to be a highly reproducible and quantitative method to determine the number of neoplastic cells in each tumor. For each mouse, we performed eight 100 μl PCR reactions per sample (4 μg DNA per reaction, 32 μg per mouse) using Q5 High-Fidelity 2x Master Mix (New England Biolabs, M0494X). The PCR products were purified with Agencourt AMPure XP beads (Beckman Coulter, A63881) using a double size selection protocol. The concentration and quality of the purified libraries were determined using the Agilent High Sensitivity DNA kit (Agilent Technologies, 5067-4626) on the Agilent 2100 Bioanalyzer (Agilent Technologies, G2939BA). The libraries were pooled based on lung weight (to have sequencing depth more evenly distributed across samples), cleaned up and size-selected using AMPure XP beads, and sequenced on the Illumina^®^HiSeq 2500 platform to generate paired-end 150 bp reads (Admera Health).

### Quantification of tumor sizes from sgID-BC sequencing data

A diagram describing the analysis steps that are part of the Tuba-seq method in more detail are summarized in **Supplemental Fig. 9a**. We used stringent filtering to identify the sgID-BC region that minimizes PCR and sequencing error, as previously described^57^. Specifically, we required no mismatch in the barcode region between the forward and reverse reads and removed all spurious tumors with the barcodes within two nucleotides from that of another larger tumor. The absolute number of cells in each tumor was calculated by scaling its sgID-BC read number with the mean read number of three spiked-in cell lines with a known absolute cell number of 5×10^5^.

### Summary statistics for tumor size distributions

As sequencing depth and PCR efficiency vary across libraries, we focused on tumors that we can repeatedly identify with high confidence, which are tumors over 500 cells as quantified by comparing technical replicates. We used multiple summary statistics to describe the truncated distribution of tumor sizes for all tumors larger than 500 cells. Percentiles and LN means were calculated as two summary statistics. Percentiles are a nonparametric summary of the distribution by taking the 50^th^, 60^th^, 70^th^, 70^th^, 90^th^, and 95^th^ percentile of the distribution. The LN mean calculates the maximum likelihood estimator of the mean tumor sizes assuming a log-normal distribution of tumor sizes. For both of these metrics, we normalized to the corresponding value of the average of inert tumors to represent the relative growth advantage of inactivating the gene.

### Quantification of treatment responses of inert tumors to osimertinib

We quantified the treatment effect of osimertinib by comparing the tumor size distributions of the vehicle- and osimertinib-treated groups. We used two ways to quantify treatment effect. The first way is to calculate the total number of neoplastic cells of tumors carrying the inert control sgRNAs in each mouse and taking the fold change of the average of the total number of neoplastic cells as an approximation of the drug effect.

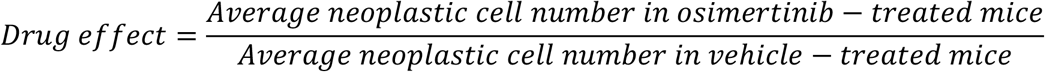

This calculation of tumor burden is very intuitive and relevant in the clinical setting, but this measure is quite variable due to the large variations in the sizes of the largest tumors. The second way assumes that each neoplastic cell, regardless of the size of the tumor harboring it, has an equal probability of being killed by osimertinib treatment (*K*). To make sure that we are evaluating tumors that are large enough to be repeatedly identified, we focus on tumors that are over 1,000 cells in the vehicle-treated mice. We estimated the tumor size reduction after treatment with osimertinib for the tumors with the inert control sgRNAs by matching the distribution of tumor sizes in vehicle- and osimertinib-treated mice. Specifically, we used the binary search algorithm to find the proportion of neoplastic cells remaining after treatment with osimertinib (*S*) between *K* = 1 (100% cells were killed by osimertinib) and *K* = 0 (0% of cells were killed by osimertinib), such that the median tumor number of tumors with the inert control sgRNAs across the vehicle-treated mice, upon simulated reduction to *K*, matches the median tumor number of tumors with the inert control sgRNAs larger than 1,000 cells across all osimertinib-treated mice.

### Estimation of genotype-specific treatment responses

We calculated the expected size distribution of tumors after treatment assuming no genotype-specific treatment responses by reducing all tumors in the vehicle-treated mice by the estimated drug effect (*K*). Then we calculated the genotype-specific treatment response for each sg*TS* by comparing the relative LN mean of all tumors in the osimertinib-treated mice and the relative LN mean of all tumors calculated from the expected distribution after treatment. The genotype-specific treatment response is calculated as the log2 ratio of the observed relative LN mean by the expected LN mean, and we named it as *ScoreRLM*. We focus on tumors with the inert control sgRNAs that are over 1,000 cells in untreated mice and take out comparable proportions of tumors with each sgRNA from each vehicle- and osimertinib-treated mice based on the estimated treatment effect and the proportion of tumors carrying each sg*TS*. The *ScoreRLM* is calculated as:

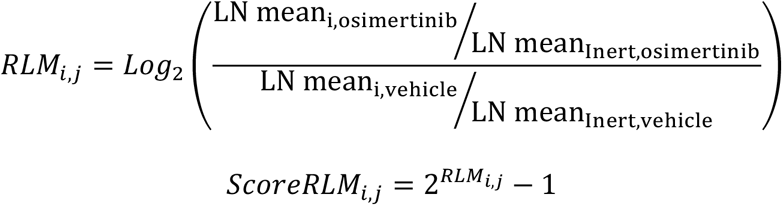

where LN mean_i,osimertinib_ is the LN mean for tumors containing sgID *i* in osimertinib-treated group, LN mean_Inert,osimertinib_ is the LN mean for all tumors containing one of the four inert sgIDs in the osimertinib-treated group. Similarly, LN mean_i,vehicle_ is the LN mean for tumors containing sgID *i* in vehicle-treated group and LN mean_Inert,vehicle_ is the LN mean for all tumors containing one of the four inert sgIDs in the vehicle-treated group. When tumors are larger than expected, the *ScoreRLM* will be positive, indicating resistance conferred by gene inactivation, while when tumors are smaller than expected, the *ScoreRLM* will be negative, indicating sensitivity conferred by tumor suppressor gene inactivation. Although the metric on the log2 scale results in the first formula with good statistical properties ranging from –∞ to + ∞ and centered on 0. To make it more directly interpretable by readers, we converted it to the linear scale as shown in the second formula. On the linear scale the metric ranges from −1 to + ∞, a value of 0.5 means the tumors are 50% larger than expected and a value of −0.5 means that the tumors are 50% smaller than expected.

Apart from *ScoreRLM* that compares the relative LN mean in the vehicle- and osimertinib-treated group, we can similarly compare the relative tumor number (*ScoreRTN*) and relative geometric mean (*ScoreRGM*) for the observed and expected distribution of tumor sizes following the same logic as shown below:

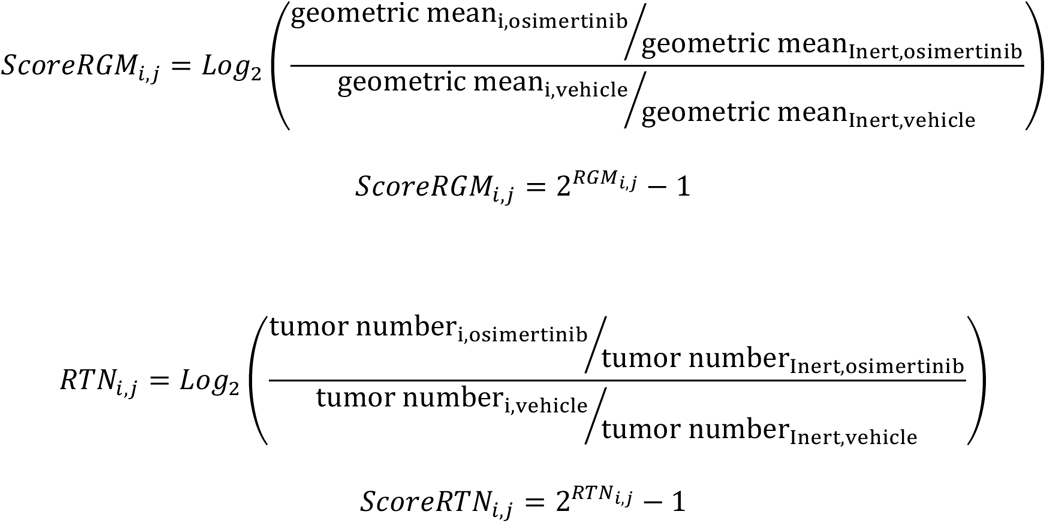

where the geometric mean and tumor number were calculated from inert tumors that are over 1,000 cells in the vehicle-treated mice and the corresponding proportions for other sgIDs in the vehicle-and osimertinib-treated groups considering the proportion of sgIDs and the treatment^57^.

The standard deviation of the genotype-specific responses, represented by any of the three metrics, is estimated by bootstrapping mice in both the vehicle- and osimertinib-treated groups and then bootstrapping tumors with the same sgID in each bootstrapped mouse. Such a two-step bootstrap process allows us to control for both variations of tumor size across mice and within the same mouse. For each run of bootstrap, we re-estimated the drug effect, estimated the expected tumor size profile between the vehicle- and osimertinib-treated mice. We estimated the standard deviation based on the scores on the log2 scale and then plotted the values on the linear scale, because the latter ranges from −1 to +∞ with no genotype-specific responses located at 1.

### Power analysis for three metrics in identifying genotype-specific treatment responses

We calculated the sensitivity and specificity for the three metrics, when 1) the genotype-specific treatment responses to osimertinib only occur in large tumors, while smaller tumors respond similarly to sg*Inert* tumors, 2) the genotype-specific treatment responses are uniform across all tumor sizes such that all sg*TS* tumors have increased or reduced sensitivity to osimertinib relative to sg*Inert* tumors (**Supplemental Figs. 6e, f**). Specifically, we used all ten vehicle-treated mice at 11 weeks for simulation. We first applied a drug effect of *K* = 75% (75% of cells are expected to be killed by osimertinib assuming no genotype-specific treatment responses) to all tumors. Then we apply the preassigned genotype-specific treatment responses to all tumors to generate the simulated distributed tumor sizes. For the first scenario of size-dependent genotype-specific treatment response, the preassigned genotype-specific responses only occur in tumors with over 10,000 cells, and they are assigned to be four-fold higher than expected, i.e., these tumors do not respond at all to osimertinib considering that 75% of tumor cells are expected to be killed without any genotype-specific treatment responses. For the second scenario of uniform responses, all tumors with the sg*TS* are assigned to be 50% larger than expected, i.e., reduced in size by 62.5% instead of 75% after treatment by osimertinib. The same sample sizes of ten vehicle- and ten osimertinib-treated mice were generated from bootstrapping mice and then bootstrapping tumors from each mouse prior to and after the simulated effects of drug responses and genotype – specific responses, respectively. To calculate the false discovery rate, we also simulated another scenario where no preassigned genotype-specific responses exist and all tumors, regardless of genotypes, were reduced in size uniformly by 75% by osimertinib. The therapeutic sensitivity was calculated as the probability of re-identifying the preassigned genotype-specific treatment responses given the cutoff of *P*-values, while specificity was calculated as the probability of falsely identifying genotype-specific treatment responses without any input signal of genotype-specific treatment responses given the cutoff of *P*-values. A total of 100 runs of simulations of the 11 tumor suppressor genes (including Tp53; 1,100 cases) were performed for each scenario. We further plotted the receiver operating characteristic^78^ curves for the three metrics by varying the cutoff for the *P*-values.

### Histology and immunohistochemistry

The auxiliary lobe of the right lung was collected for each experimental mouse from the mice transduced with the Lenti-sg*TS*^*Pool*^/*Cre* pool whereas, both the auxiliary and left lobes were collected for the validation experiments with individual sgRNAs (sg*Apc*, sg*Rbm10*, sg*Rbm10#2* and sg*Neo2*). Right and auxiliary lobes were collected for the experiment with Lenti-sg*Keap1/Cre* virus. Lung lobes were fixed in 4% paraformaldehyde overnight at room temperature, placed in 70% ethanol, and paraffin-embedded and sectioned (Histology @ Yale). Four micrometer sections were used for hematoxylin and eosin (H&E) staining and immunohistochemistry. Tumor sizes were determined by measuring the longest diameter for each tumor in H&E stained sections. Tumor size and tumor area were quantified using ImageJ. The limited tissue collected for histological analysis reduced the number of tumors initiated with Lenti-sg*Rbm10/Cre* that could be measured. *P*-values were calculated from the Mann-Whitney U test. The following antibodies were used for immunohistochemistry: anti-mutant EGFR^L858R^ (1:200, CST #3197), anti-SP-C (1:200, AB40876), anti-TTF-1/Nkx2-1 (1:200, AB76013), anti-phospho-histone H3 (1:200, CST #9701), anti-Ki-67 (1:400, CST #9027), anti-mKate (1:2500, Evrogen #AB233), anti-Cas9 (1:500, Novus Biologicals #7A9-3A3, NBP2-36440), anti-β-catenin (1:500, CST #8814), anti-Rbm10 (1:200, AB224149), and anti-Keap1 (1:500, AB227828). Rbm10 expression in tumors was binned as High (over 75% of positive nuclei), Medium (between 25 and 75% of positive nuclei) or Low/negative (positive nuclear staining below 25%). Keap1 levels in tumors were binned as High (over 75% of positive cells), Medium (between 25 and 75% of positive cells) or Low/negative (positive staining below 25%). *P*-values were calculated using the Chi-squared test.

### Analysis of human lung tumor data using GENIE data

The AACR Project GENIE is a registry that contains CLIA-/ISO-certified genomic data collected from the records of more than 9,000 patients who were treated at each of the consortium’s participating institutions^48^. Data from GENIE version 4.1-public were accessed through the Synapse Platform. We generated a comprehensive list of all missense, nonsense, and frameshift mutations for all screened genes across all participating centers in the GENIE project, documenting these mutations at the gene and amino acid levels. Based on the sets of genes included in the different screening panels that contribute to Project GENIE, we annotated all lung adenocarcinoma within the database as being wild-type, mutated, or not screened for each gene. From this complete catalog of mutations in each tumor, we determined the rates of co-occurrence of known oncogenic *KRAS* mutations (G12X, G13X, Q61X) or *EGFR* mutations (L858R, Exon 19 deletions, L861Q, G719X) with missense, nonsense, or frameshift mutations in our set of ten tumor suppressor genes. Furthermore, we determined the frequency of co-incident tumor suppressor gene mutations in tumors with oncogenic *KRAS* and inactivating *TP53* mutations (*KRAS/TP53*) and in tumors with oncogenic *EGFR* and inactivating *TP53* mutations (*EGFR/TP53*). A similar analysis was also performed on tumors with *ALK*-rearrangements and *TP53* mutations using GENIE version 7.0 (**Supplemental Fig. 11**). A pipeline for this analysis is publicly available at www.github.com/dgmaghini/GENIE, which allows users to input OncoTree codes of interest, generate mutational profiles for the corresponding GENIE tumors, and identify the co-occurrence of up to two background mutations (for example, *EGFR* and *TP53*) with mutations in other genes of interest.

### Stanford cohort of *EGFR* mutant lung adenocarcinomas

Patients with lung cancer who were evaluated at the Stanford Cancer Center and had their tumors analyzed using the Stanford Solid Tumor Actionable Mutation Panel (STAMP)^6^ were included in the analysis. This retrospective study was conducted under a molecular analysis protocol approved by the Stanford University Institutional Review Board. All STAMP cases performed between 2015 and 2019 were included; during this time, there were two different assays used, with one covering 198 genes (302 kb) and the other covering 130 genes (232 kb). STAMP was done as the standard of care and thus at the discretion of the treating physician and could have occurred at the time of diagnosis or at the time of progression.

From the available STAMP cases, patients were selected if they had stage IV non-small cell lung cancer (NSCLC) and had a pathogenic *EGFR* and *p53* mutation. Patients were excluded if they had incomplete data or were lost to follow up prior to analysis of the primary endpoint, if they elected not to receive treatment or if they received adjuvant tyrosine kinase inhibitor therapy for early-stage disease. Within this cohort, patients were further selected for the presence of at least one of the tumor suppressor genes investigated in the preclinical setting (*KEAP1, LKB1/STK11, SETD2, SMAD4, RB1, APC, ARIDIA* and/or *CDKN2A*). To ensure that there was enough power to conduct an analysis, genes were analyzed individually if there were at least 8 patients with tumors with a mutation of that gene who met inclusion criteria. Due to small numbers, patients with tumors with *LKB1* (*N* = 1), *SETD2* (*N* = 0) and *SMAD4* (*N* = 1) mutations were not analyzed separately. However, as the two patients with tumors harboring *LKB1* and *SMAD4* mutations were also *EGFR/P53* mutant, they were included as part of the control arm. Univariate analysis of time to treatment failure was completed on cases with stage IV, *EGFR/P53* mutant NSCLC who were treated with a tyrosine kinase inhibitor ^13^ and had mutations in either *KEAP1-*pathway component *(KEAP1, NFE2L2, CUL3), RB1, APC, ARIDIA* or *CDKN2A.* Mutations that were significant on univariate analysis were identified and multivariate analyses accounting for 1) co-founding variables and 2) co-mutations were run. Demographic data including sex, age at diagnosis, smoking history, and ethnicity were extracted (**Supplemental Tables 1, 2**). For each tumor suppressor, patients were matched 1:3 with a wildtype cohort on the basis of sex, smoking history, ethnicity, age and treatment type. Time to Treatment Failure ^79^ was determined by subtracting the date of discontinuation of TKI due to progression, toxicity or death, from the date of initiation of TKI and reported in months. Statistical analysis was performed using Prism 8 and R studio. The Kaplan-Meier method was used to estimate TTF. Comparison of survival curves was made using the Log-rank test. Significance was defined as *P* < 0.05. Hazard ratios (HR) were generated from multivariate regression analysis performed in R and reported with a 95% confidence interval (CI).

### Yale cohort of *EGFR* mutant lung adenocarcinomas

Patients with *EGFR*-mutant lung adenocarcinoma were consented and enrolled to a Yale University IRB approved protocol allowing the collection and analysis of clinical data, archival and fresh tissue, blood and the generation of patient-derived xenografts. Patients who received targeted therapy (erlotinib, gefitinib, or afatinib) as first line therapy, either alone or in combination with other therapies such as chemotherapy or cetuximab, were included (**Supplemental Table 3**). For genomic studies, formalin-fixed paraffin embedded tissue was macro-dissected to enrich for tumor material. All of the tumor samples (*N* = 29) analyzed had matched normal tissue and cancer cell purity >20% as assessed by the LOH (Loss of Heterozygosity) events from the whole exome sequencing data.

#### Whole Exome Sequencing

DNA was extracted from and analyzed as previously described^80^. Briefly, genomic DNA was captured on the NimbleGen 2.1M human exome array and subjected to 74-bp paired-end reads on the Illumina HiSeq2000 instrument. The mean coverage for normal was 109.1x and the mean coverage for tumor was 189x with 92.78% and 95.56% for the bases covered at least 20 independent sequence reads, respectively. Sequence reads were mapped to the human reference genome (GRCh37) using the Burrow-Wheeler Aligner-MEM (BWA-MEM) program. Sequence reads outside the targeted sequences were discarded and the statistics on coverage were collected from the remaining reads using in-house per scripts.

#### Somatic Mutation Calling

For all matched tumor-normal pairs, somatic point mutations and indels were called by MuTect2 using Bayesian classifiers. For all somatic mutations called, we extracted base coverage information in all samples and considered the mutations that were supported by at least two independent sequence reads covering non-reference alleles and present in more than 5% of all sequencing reads. Identified variants were further filtered based on their presence in repositories of common variations (1000 Genomes, NHLBI exome variant server and 2,577 non-cancer exomes sequenced at Yale) and annotated using ANNOVAR program^81^. All somatic indels were visually inspected to remove the false positive calls.

#### Somatic Copy Number Variation Analysis

Copy number analysis was performed as previously described^80^. Briefly, copy number variants were identified from the whole exome sequencing data using EXCAVATOR software that normalizes the non-uniform whole exome sequencing data taking GC-content, mappability, and exon-size into account^82^. The Hidden Markov Model was utilized to classify each copy number variant segment into five copy number states (homozygous deletion, heterozygous deletion, normal copy number, homozygous copy gain or multiple copy gain). Tumor purity was estimated from LOH using in-house per scripts.

## Data availability

The datasets generated during and/or analyzed in the current study will be made available in the NCBI Gene Expression Omnibus database (GSE146550) and dbGAP. GENIE genomic data analysis is publicly available at www.github.com/dgmaghini/GENIE. All other data supporting the findings are available upon request.

## Code availability

The code is available at https://github.com/lichuan199010/Tuba-seq-analysis-and-summary-statistics.

