## Supplemental Data for "Genetic determinants of *EGFR*-Driven Lung Cancer Growth and Therapeutic Response *In Vivo*"

**a**

| | | Tumor context | |
|---|---|---|---|
| ***TS*** | **KO system** | **In oncogenic *Kras*-driven models** | **In oncogenic *EGFR*-driven models** |
| **Pool** | | | |
| *Apc* | CRISPR | Sánchez-Rivera *et al.*, 2014; Rogers *et al.*, 2017; Rogers *et al.*, 2018 | No published data |
| *Arid1a* | CRISPR | Walter *et al.*, 2017; Rogers *et al.*, 2017; Rogers *et al.*, 2018 | |
| *Atm* | Null alleles; Cre/loxP; CRISPR | Efeyan *et al.*, 2009; Schmitt et al., 2017; Rogers *et al.*, 2017; Rogers *et al.*, 2018 | |
| *Cdkn2a* | Null alleles; Cre/loxP; CRISPR | Fisher et al., 2001; Schuster *et al.*, 2014; Rogers *et al.*, 2017; Rogers *et al.*, 2018 | |
| *Keap1* | CRISPR | Romero *et al.*, 2017; Rogers *et al.*, 2017; Rogers *et al.*, 2018 | |
| *Rb1* | Cre/loxP; CRISPR | Ho *et al.*, 2009; Rogers *et al.*, 2017; Rogers *et al.*, 2018; Walter *et al.*, 2019 | |
| *Rbm10* | CRISPR | Rogers *et al.*, 2017; Rogers *et al.*, 2018 | |
| *Setd2* | CRISPR | Walter *et al.*, 2017; Rogers *et al.*, 2017; Rogers *et al.*, 2018 | |
| *Smad4* | CRISPR | Rogers *et al.*, 2017; Rogers *et al.*, 2018 | |
| *Lkb1* | Cre/loxP; CRISPR | Ji *et al.*, 2007; Rogers *et al.*, 2017; Rogers *et al.*, 2018 | |
| *Tp53* | Cre/loxP; CRISPR | Jackson *et al.*, 2005; Rogers *et al.*, 2017; Rogers *et al.*, 2018 | |
| **Others** | | | |
| *Cdkn2a/b* | Cre/loxP | Schuster *et al.*, 2014 | No published data |
| *Dnmt3a* | Cre/loxP | Goa *et al.*, 2011 | |
| *Kdm6a* | Cre/loxP | Wu *et al.*, 2018 | |
| *Nf1* | CRISPR | Wang *et al.*, 2019 | |
| *Pten* | Cre/loxP; CRISPR | Iwanaga *et al.*, 2008; Curry *et al.*, 2013; Sánchez-Rivera *et al.*, 2014 | |
| *Smarca4* | CRISPR | Walter *et al.*, 2017 | |
| *Tsc1* | Cre/loxP | Liang *et al.*, 2010 | |
| *Notch2* | Cre/loxP | Baumgart *et al.*, 2014 | |
| *Mig6* | Null alleles | | Maity *et al.*, 2015 |

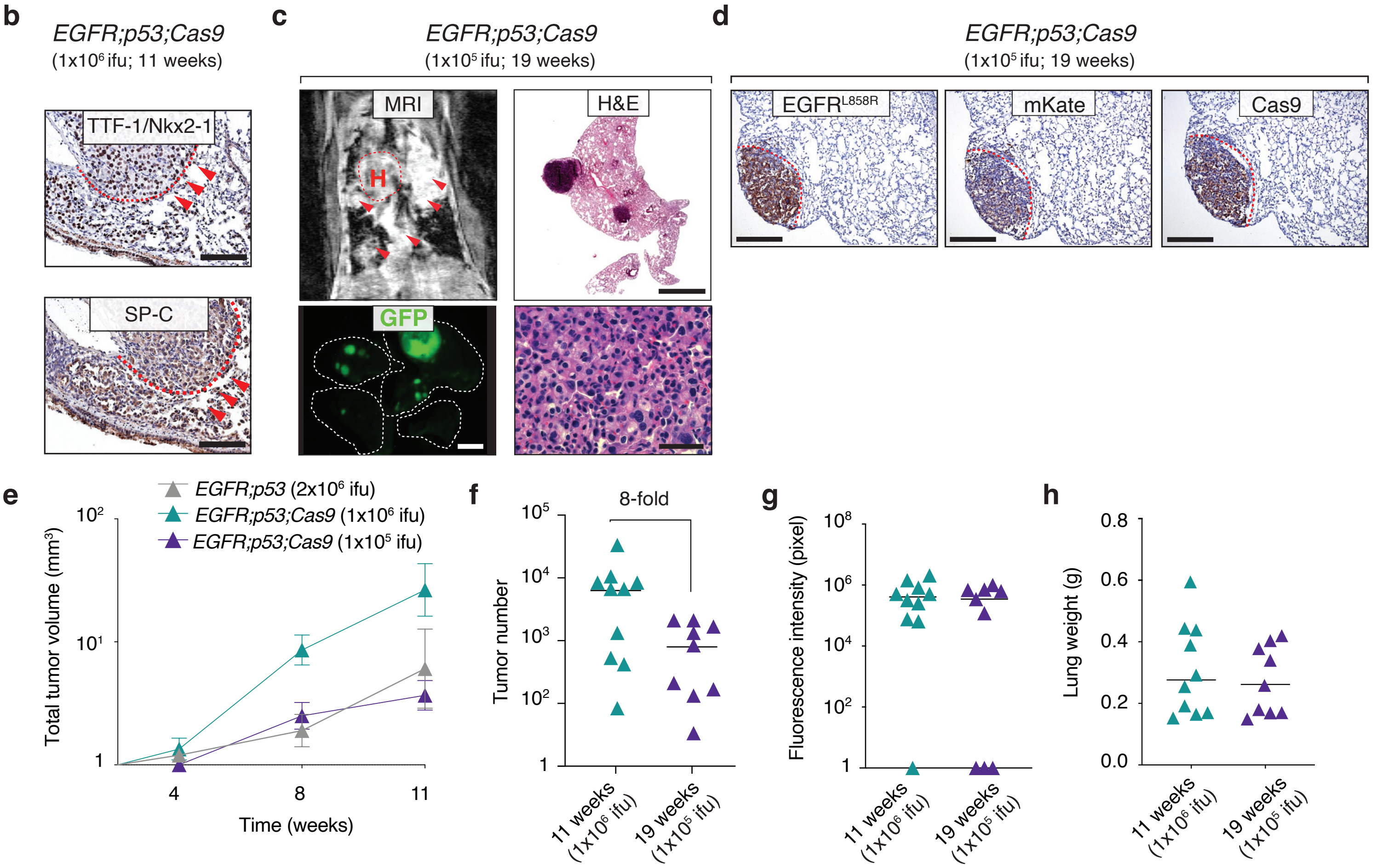


**Supplemental Fig. 1 | Characteristics of lung tumors in *EGFR;p53;Cas9* mice. a**, Summary of the studies on tumor suppressor gene inactivation *in vivo*. Very few published data are currently available for the *EGFR*-driven lung cancer model. **b**, Immunohistochemical staining showing a tumor positive for TTF-1/Nkx2-1 and SP-C in *EGFR;p53* mice. Red arrows indicate tumor areas. Scale bars = 100 µm. **c**, MRI showing tumor development 19 weeks after tumor initiation in a representative *EGFR;p53;Cas9* mouse ($N = 9$) following delivery with $1x10^5$ ifu of Lenti-sg*TS*$^{Pool}$/*Cre* virus. H&E staining showing lung adenocarcinomas (right panels). Scale bars = 1.2 mm and 100 µm for the top and bottom panel, respectively. The dashed lines indicate the lungs. GFP image scale bar = 2.5 mm. **d**, Immunohistochemical staining for EGFR$^{L858R}$, mKate and Cas9 after 19 weeks of tumor initiation in *EGFR;p53;Cas9* mice. The red lines indicate tumor areas. Scale bars = 200 µm. **e**, Tumor growth curves in *EGFR;p53* and *EGFR;p53;Cas9* mice transduced with Lenti-sg*TS*$^{Pool}$/*Cre* virus quantified by measuring tumor volume by MRI at the indicated time points. Each triangle indicates the average of all tumor volumes from each group. Standard errors of the mean are shown. **f**, Comparison of tumor number determined by Tuba-seq in *EGFR;p53;Cas9* mice transduced with Lenti-sg*TS*$^{Pool}$/*Cre* virus ($1x10^6$ versus $1x10^5$ ifu). Tumors initiated with a lower titer for 19 weeks are bigger and significantly fewer compared to tumors transduced with $1x10^6$ ifu of Lenti-sg*TS*$^{Pool}$/*Cre* virus. Each triangle represents a mouse. Horizontal lines show the median. **g**, GFP quantification of tumor burden in *EGFR;p53;Cas9* mice after 11 or 19 weeks of tumor initiation. Horizontal lines show the median. **h**, Tumor burden represented as the weight of tumor-bearing lungs in *EGFR;p53;Cas9* mice after 11 and 19 weeks of tumor initiation. Horizontal lines indicate the median.

**a**

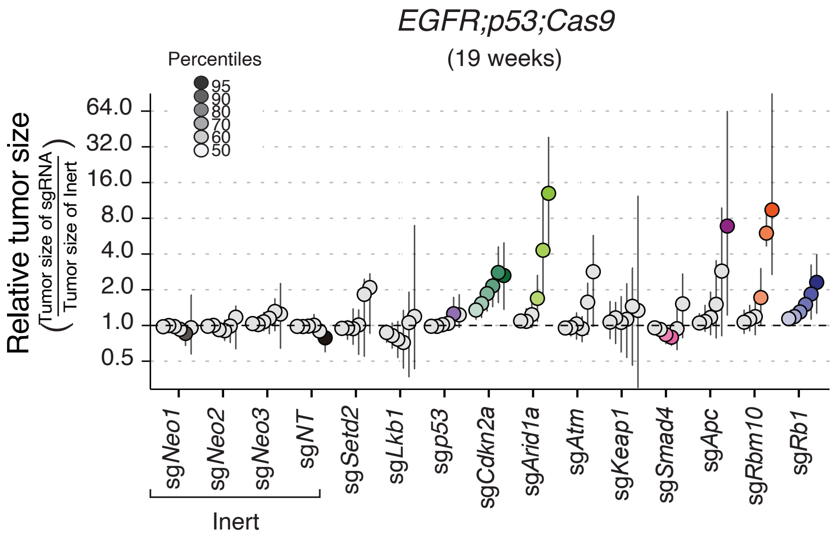


**b**

*EGFR;p53;Cas9*

| | | mean | *P*-value |
|---|---|---|---|
| | sg*Rb1* | **1.71** | **0.03** |
| | sg*Rbm10* | **4.41** | **0.0002** |
| | sg*Apc* | 4.77 | 0.06 |
| | sg*Smad4* | 0.95 | 0.78 |
| | sg*Keap1* | 1.14 | 0.80 |
| | sg*Atm* | 1.45 | 0.31 |
| | sg*Arid1a* | **3.24** | **0.003** |
| | sg*Cdkn2a* | **1.82** | **0.01** |
| | sg*p53* | 1.08 | 0.58 |
| | sg*Lkb1* | 0.86 | 0.62 |
| | sg*Setd2* | 1.26 | 0.40 |
| Inert | sg*Neo1* | 1.04 | 0.82 |
| Inert | sg*Neo2* | 1.00 | 0.97 |
| Inert | sg*Neo3* | 1.03 | 0.87 |
| Inert | sg*NT* | 0.94 | 0.41 |

**c** **d**

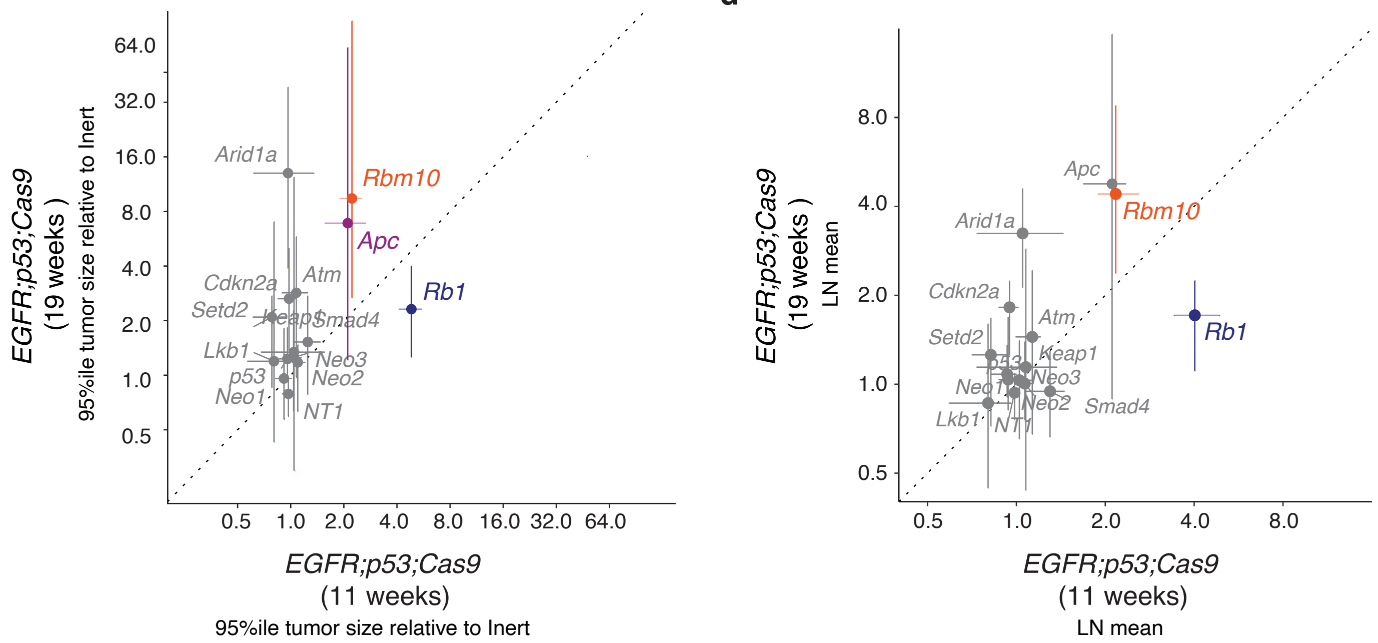


**Supplemental Fig. 2 I Tuba-seq uncovers positive and negative effects of putative tumor suppressor gene inactivation on *EGFR*-driven lung tumor growth. a**, Relative size of tumors of each genotype in *EGFR;p53;Cas9* mice (*N* = 9) 19 weeks after tumor initiation with Lenti-sg*TS*$^{Pool}$/*Cre* virus. *P*-values are calculated from bootstrapping. Percentiles that are significantly different from the tumors with inert sgRNAs are in color. 95% confidence intervals are shown. **b**, LN mean for tumors with each sgRNA in *EGFR;p53* and *EGFR;p53;Cas9* mice 19 weeks after tumor initiation (normalized to the tumors with inert sgRNAs). *P*-values are calculated from bootstrapping. *P*-values < 0.05 and their corresponding means are highlighted bold when the size effects are equal to or >10% compared to the size of tumors with inert sgRNAs. **c**, Comparison of the relative effect of tumor suppressor gene inactivation in *EGFR;p53;Cas9* mice after 11 (X-axis) and 19 weeks (Y-axis) after tumor initiation. Relative tumor size for each sg*TS* compared to inert at the 95th percentile 11 and 19 weeks after tumor initiation with 1x10$^{6}$ and 1x10$^{5}$ ifu of Lenti-sg*TS*$^{Pool}$/*Cre* virus respectively (*EGFR;p53;Cas9* mice, 11 weeks *N* = 10; *EGFR;p53;Cas9* mice, 19 weeks *N* = 9). Error bars show the 95% confidence intervals. Genotypes that are significantly different from the inert tumors for both time-points are highlighted in color. **d**, Relative LN mean comparison of tumor suppressor gene inactivation in *EGFR;p53;Cas9* after 11 and 19 weeks after tumor initiation. 95% confidence intervals are shown. Only genotypes that are significantly different from the inert for both time points, are highlighted in color.

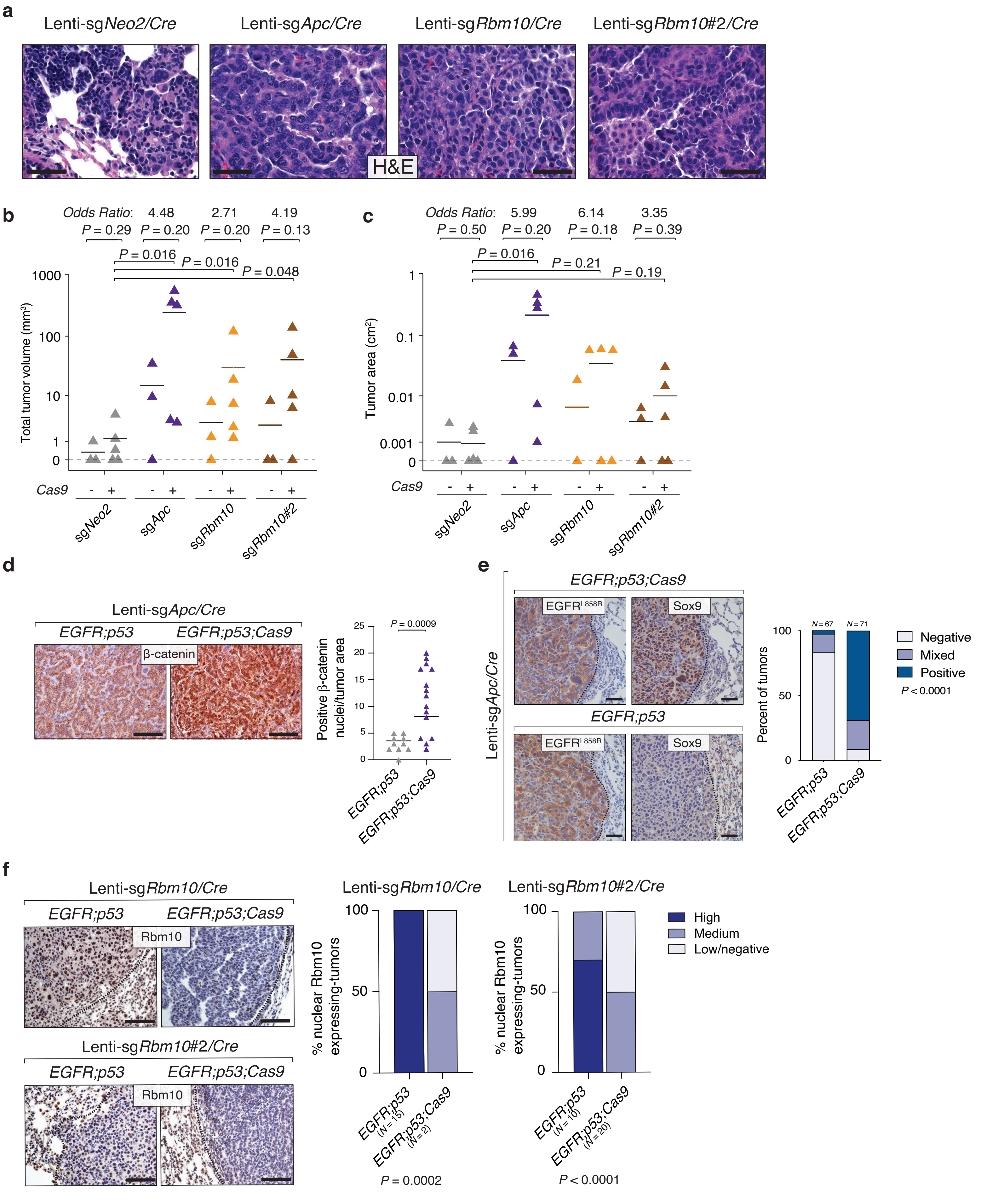


**Supplemental Fig. 3 | *Apc* and *Rbm10* inactivation with individual Lenti-sg*TS/Cre* viruses confirm Tuba-seq results.**

**a**, H&E images showing lung adenocarcinomas in *EGFR;p53;Cas9* mice after 14 weeks of tumor initiation with individual Lenti-sg*TS/Cre* (sg*Apc*, sg*Rbm10,* sg*Rbm10#*2 and sg*Neo2*). Scale bars = 100 µm. **b**, Total tumor burden 14 weeks after tumor initiation as quantified by MRI in *EGFR;p53* and *EGFR;p53;Cas9* mice in which tumors were initiated with the indicated vectors. Horizontal lines indicate the median. *P*-values were calculated using a one-tailed Mann-Whitney U test. The odds ratio is a measure of the magnitude of increase in tumor volume conferred by inactivation of the gene of interest. **c**,Tumor burden after 14 weeks of tumor initiation as quantified by total tumor area in *EGFR;p53;Cas9* mice transduced with individual Lenti-sg*TS/Cre* viruses compared to *EGFR;p53* mice and *EGFR;p53;Cas9* mice transduced with Lenti-sg*Neo2/Cre* virus. Tumor area was calculated from one tumor-bearing lung section. *P*-values were calculated using a one-tailed Mann-Whitney U test. The odds ratios are a measure of the strength of the increase in tumor area conferred by the corresponding gene inactivation. Horizontal lines indicate the median. **d**, Immunostaining and quantification of nuclear β-catenin localization in *EGFR;p53;Cas9* mice transduced with Lenti-sg*Apc/Cre* compared to *EGFR;p53* mice. Scale bars = 100 µm. Each triangle represents positive nuclear staining for each tumor area. Tumor area corresponds to a tumor-bearing lung histological field ($N = 10$ in *EGFR;p53* mice and $N = 15$ in *EGFR;p53;Cas9* mice). Horizontal lines indicate the median. The *P*-value was calculated using the Mann-Whitney U test. **e**, Immunostaining and quantification of nuclear Sox9 in *EGFR;p53;Cas9* compared to *EGFR;p53* tumors (as Positive, Mixed or Negative) initiated with Lenti-sg*Apc/Cre*. The dashed lines indicate areas of tumors. Scale bars = 50 µm. *P*-values were calculated using a Chi-squared test. **f**, Immunostaining and evaluation of Rbm10 expression (high, medium or low/negative) in tumors initiated in *EGFR;p53;Cas9* mice with either Lenti-sg*Rbm10/Cre* (top panels) or Lenti-sg*Rbm10#2/Cre* (bottom panels) compared to *EGFR;p53* mice (left panels). The dashed lines indicate areas of tumors. Scale bars = 100 µm. *P*-values were calculated using a Chi-squared test.

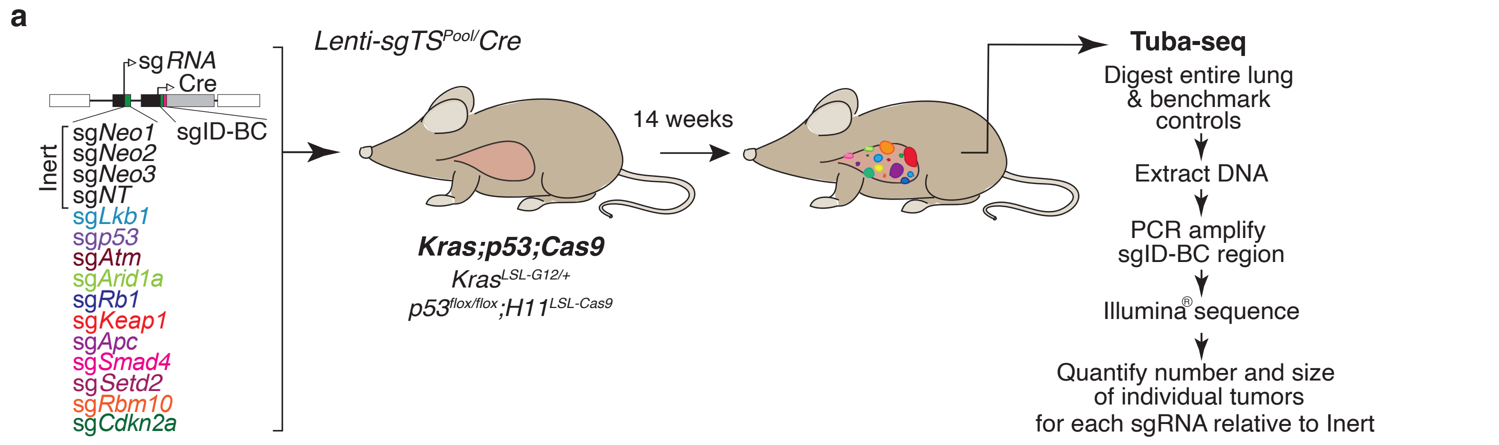


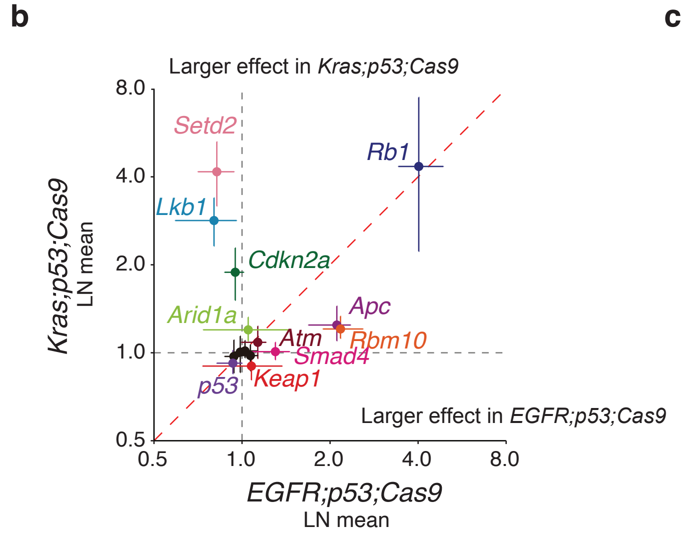


**c**

| | *LKB1* wt | *LKB1* mut | |
|---|---|---|---|
| *EGFR^onc* | 542 | 8 | 1.5% |
| *KRAS^onc* | 1436 | 319 | 18.2% |

Odds ratio = 15.05
*P* = $1.7 \times 10^{-30}$

| | *RBM10* wt | *RBM10* mut | |
|---|---|---|---|
| *EGFR^onc* | 250 | 33 | 11.7% |
| *KRAS^onc* | 621 | 81 | 11.5% |

Odds ratio = 0.99
*P* = 1.00

| | *SETD2* wt | *SETD2* mut | |
|---|---|---|---|
| *EGFR^onc* | 390 | 13 | 3.2% |
| *KRAS^onc* | 1148 | 75 | 6.1% |

Odds ratio = 1.96
*P* = 0.03

| | *APC* wt | *APC* mut | |
|---|---|---|---|
| *EGFR^onc* | 527 | 23 | 4.2% |
| *KRAS^onc* | 1680 | 75 | 4.3% |

Odds ratio = 1.02
*P* = 1.00

**Supplemental Fig. 4 I Oncogenic drivers dictate the fitness landscape of tumor suppression. a**, Tumors were initiated in *Kras;p53;Cas9* mice with Lenti-sg*TS^Pool/Cre* virus ($2.2 \times 10^4$ ifu) and collected for Tuba-seq 14 weeks after tumor initiation (*N* = 6). **b**, LN mean for different tumor suppressor genes in *EGFR;p53;Cas9* and *Kras;p53;Cas9* mice. *P*-values were calculated from bootstrapping. Confidence intervals are shown. **c**, Frequency of co-occurrence of *LKB1, RBM10, SETD2* and *APC* alterations with *EGFR^onc* and *KRAS^onc* alterations in human lung adenocarcinomas. The odds ratios represent a measure of the strength of the frequency of co-occurring tumor suppressor gene alterations with *KRAS^onc* compared to *EGFR^onc* cases. *P*-values were calculated using a Fisher's exact test.

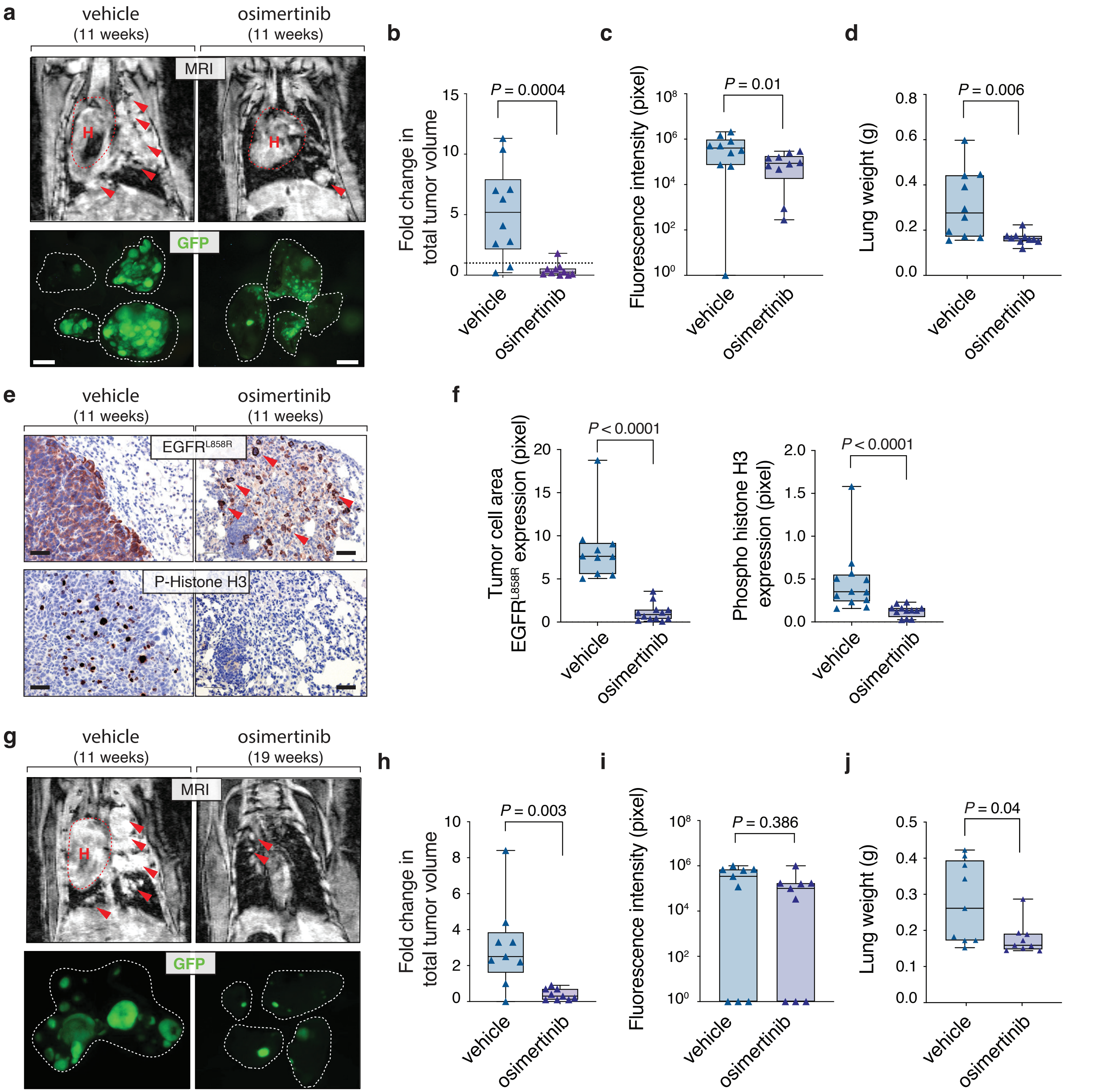


**Supplemental Fig. 5 l Strong therapeutic response to the 3rd generation TKI, osimertinib at 11 and 19 weeks after tumor initiation. a**, *EGFR*-driven lung adenocarcinomas and early-therapeutic response to osimertinib. Tumor burden visualized using MRI and GFP fluorescence after two weeks of osimertinib treatment (25 mg/kg 5 days/week) at 11 weeks after tumor initiation compared to vehicle. Scale bars = 2.5 mm. **b**-**d**, Overall response to osimertinib treatment at 11 weeks of tumor initiation. Therapeutic response is measured as (**b**) the fold change in total tumor volume relative to baseline (by MRI quantification), (**c**) change in GFP fluorescence, and (**d**) weight of tumor-bearing lungs after treatment with vehicle or osimertinib. *P*-values were calculated using the Mann-Whitney U test. Horizontal lines show the median. **e**, Immunostaining and relative quantification of mutant EGFR positive cells and phospho-histone H3 positive tumor cells showing that tumors treated with osimertinib are less proliferative compared to tumors treated with vehicle. Red arrows indicate residual tumor cells. Scale bars = 100 μm. **f**, Quantification of mutant EGFR positive and phospho-histone H3 positive tumor cells in vehicle- and osimertinib-treated tumors after 11 weeks of tumor initiation. Each triangle represents a tumor area in a histological lung section. *P*-values were calculated using the Mann-Whitney U test. Horizontal lines indicate the median. **g**, Tumor burden visualized by MRI and GFP fluorescence after two weeks of treatment with vehicle or osimertinib 17 weeks after tumor initiation. **h**-**j**, Overall response to osimertinib at 19 weeks of tumor initiation measured as (**h**) the fold change in total tumor volume relative to baseline, (**i**) change in GFP fluorescence and (**j**) a change in the weight of tumor-bearing lungs. *P*-values were calculated using the Mann-Whitney U test. Horizontal lines indicate the median.

**a**

*EGFR;p53;Cas9* + osimertinib

| | (11 weeks) LN mean | (11 weeks) *P*-value | (19 weeks) LN mean | (19 weeks) *P*-value |
|---|---|---|---|---|
| sg*Rb1* | **1.87** | **<10⁻⁴** | 1.02 | 0.73 |
| sg*Rbm10* | **1.35** | **<10⁻⁴** | **3.08** | **<10⁻⁴** |
| sg*Apc* | **1.49** | **<10⁻⁴** | **2.08** | **0.001** |
| sg*Smad4* | 1.03 | 0.35 | 0.81 | 0.05 |
| sg*Keap1* | **1.41** | **<10⁻⁴** | **2.06** | **0.04** |
| sg*Atm* | 0.98 | 0.51 | 0.95 | 0.50 |
| sg*Arid1a* | 0.95 | 0.04 | 1.34 | 0.23 |
| sg*Cdkn2a* | 1.03 | 0.15 | **1.29** | **0.003** |
| sg*p53* | 1.00 | 0.85 | 0.90 | 0.67 |
| sg*Lkb1* | 0.90 | 0.01 | 0.92 | 0.81 |
| sg*Setd2* | 0.95 | 0.05 | 1.37 | 0.26 |
| Inert: sg*Neo1* | 1.01 | 0.74 | 0.89 | 0.05 |
| Inert: sg*Neo2* | 1.04 | 0.31 | 1.16 | 0.54 |
| Inert: sg*Neo3* | 0.99 | 0.66 | **1.18** | **<10⁻⁴** |
| Inert: sg*NT* | 0.98 | 0.29 | 0.87 | 0.21 |

**b**

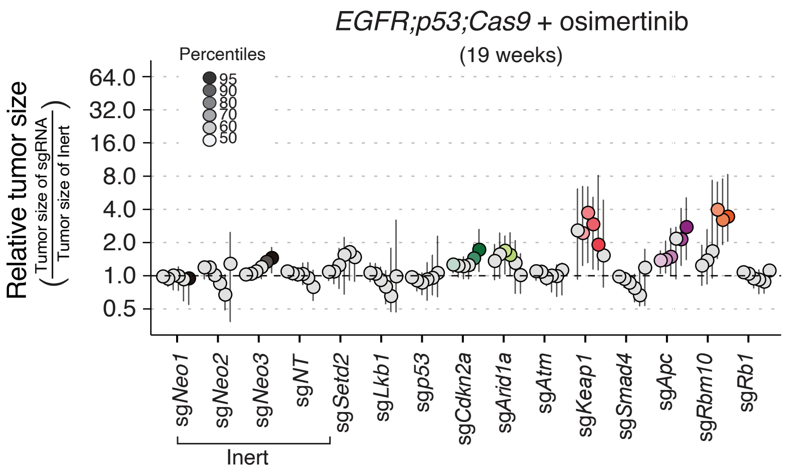


**c**

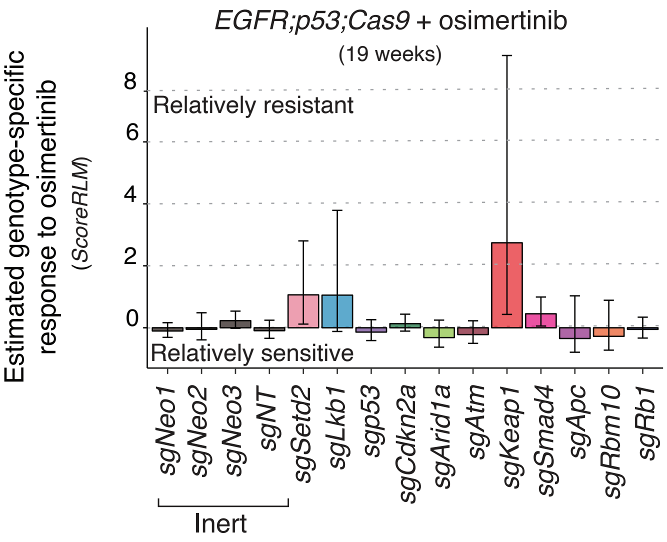


**d**

*EGFR;p53;Cas9* + osimertinib (11 weeks)

| | *ScoreRGM* Score | *ScoreRGM* *P*-value | *ScoreRTN* Score | *ScoreRTN* *P*-value |
|---|---|---|---|---|
| sg*Rb1* | -0.38 | 0.06 | 0.01 | 0.37 |
| sg*Rbm10* | -0.15 | 0.33 | 0.12 | 0.20 |
| sg*Apc* | -0.16 | 0.23 | -0.03 | 0.56 |
| sg*Smad4* | 0.10 | 0.39 | 0.26 | 0.09 |
| sg*Keap1* | 0.37 | 0.28 | -0.02 | 0.64 |
| sg*Atm* | -0.12 | 0.21 | -0.06 | 0.31 |
| sg*Arid1a* | 0.02 | 0.58 | -0.02 | 0.57 |
| sg*Cdkn2a* | -0.02 | 0.48 | -0.08 | 0.39 |
| sg*p53* | 0.06 | 0.30 | 0.03 | 0.43 |
| sg*Lkb1* | 0.18 | 0.31 | 0.30 | 0.32 |
| sg*Setd2* | 0.03 | 0.39 | 0.03 | 0.39 |
| Inert: sg*Neo1* | 0.02 | 0.41 | 0.01 | 0.50 |
| Inert: sg*Neo2* | -0.02 | 0.47 | 0.00 | 0.64 |
| Inert: sg*Neo3* | 0.02 | 0.49 | 0.02 | 0.40 |
| Inert: sg*NT* | -0.03 | 0.41 | -0.02 | 0.40 |

**e**

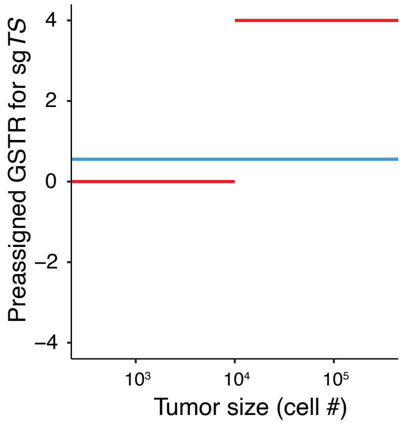


**f**

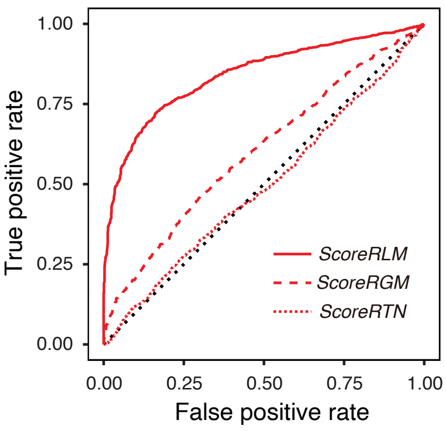


**g**

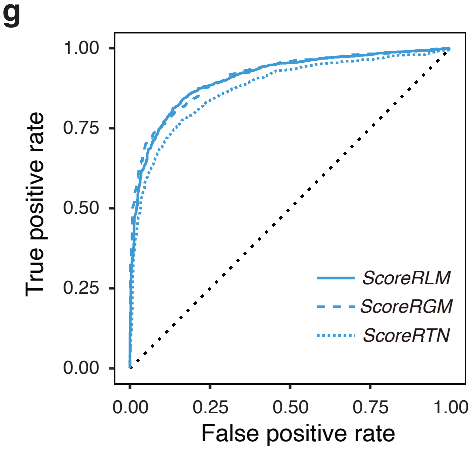


**Supplemental Fig. 6 | *Keap1* inactivation limits sensitivity to treatment with osimertinib.**

**a**, LN mean of osimertinib-treated tumors with each sgRNA normalized to inert tumors in *EGFR;p53;Cas9* mice at 11 and 19 weeks after tumor initiation. *P*-values were calculated from bootstrapping. *P*-values < 0.05 and their corresponding means are highlighted in red or green when the size effects are equal to or >10% compared to the size of tumors with inert sgRNAs. **b**, Graph of the relative size of tumors of each genotype in *EGFR;p53;Cas9* mice (*N* = 9) 19 weeks after tumor initiation and two weeks of osimertinib treatment. The relative size of tumors at the indicated percentiles was calculated from the tumor size distribution of all tumors from nine mice. Error bars show the 95% confidence intervals. *P*-values were calculated from bootstrapping. Percentiles that are significantly different from the tumors with inert sgRNAs are in color. **c**, Estimate of the genotype-specific treatment response (*ScoreRLM*) calculated by comparing the LN mean of tumors treated with osimertinib to the LN mean of vehicle-treated tumors in *EGFR;p53;Cas9* mice at 19 weeks of tumor initiation. Error bars indicate standard deviations. *P*-values were calculated from bootstrapping. None were significant, although *Keap1* inactivation is associated with a large magnitude of effect for decreased sensitivity. **d**, Tabulation of two other metrics for genotype-specific responses based on the geometric mean (*ScoreRGM*) and the relative tumor number (*ScoreRTN*) do not show significant genotype-specific effects. *P*-values were calculated from bootstrapping. **e**-**g**, Power analysis of each summary statistics (*ScoreRGM*, *ScoreRTN* and *ScoreRLM*) to uncover genotype-specific treatment responses (GSTRs). To generate tumor size profiles with known GSTRs, we generated simulated data from bootstrap resampling of *EGFR;p53;Cas9* vehicle-treated mice 11 weeks after tumor initiation (*N* = 10, from Fig. 2). We simulated an overall drug effect by reducing the size of all tumors in the simulated drug-treated group by 75%. **e,** The red and blue lines show two examples of preassigned potential GSTRs on sgTS tumors. The red line represents a case where sg*TS* tumors larger than 10,000 cells do not respond to the drug at all, thus being 4-fold larger than expected without a GSTR (only the largest tumors show drug resistance). The blue line represents a case where all sg*TS* tumor were reduced in size by only 50%, thus being 50% larger than expected without a GSTR (equal resistance across all tumor sizes). **f,** When identifying GSTRs as shown in the red lines in (**e**), *ScoreRTN* and *ScoreRGM* were much less sensitive than *ScoreRLM*. **g,** When identifying GSTRs as shown in the blue line in (**e**), all three metrics showed reasonable and similar statistical power. Therefore, we used *ScoreRLM* as the summary statistic throughout this manuscript.

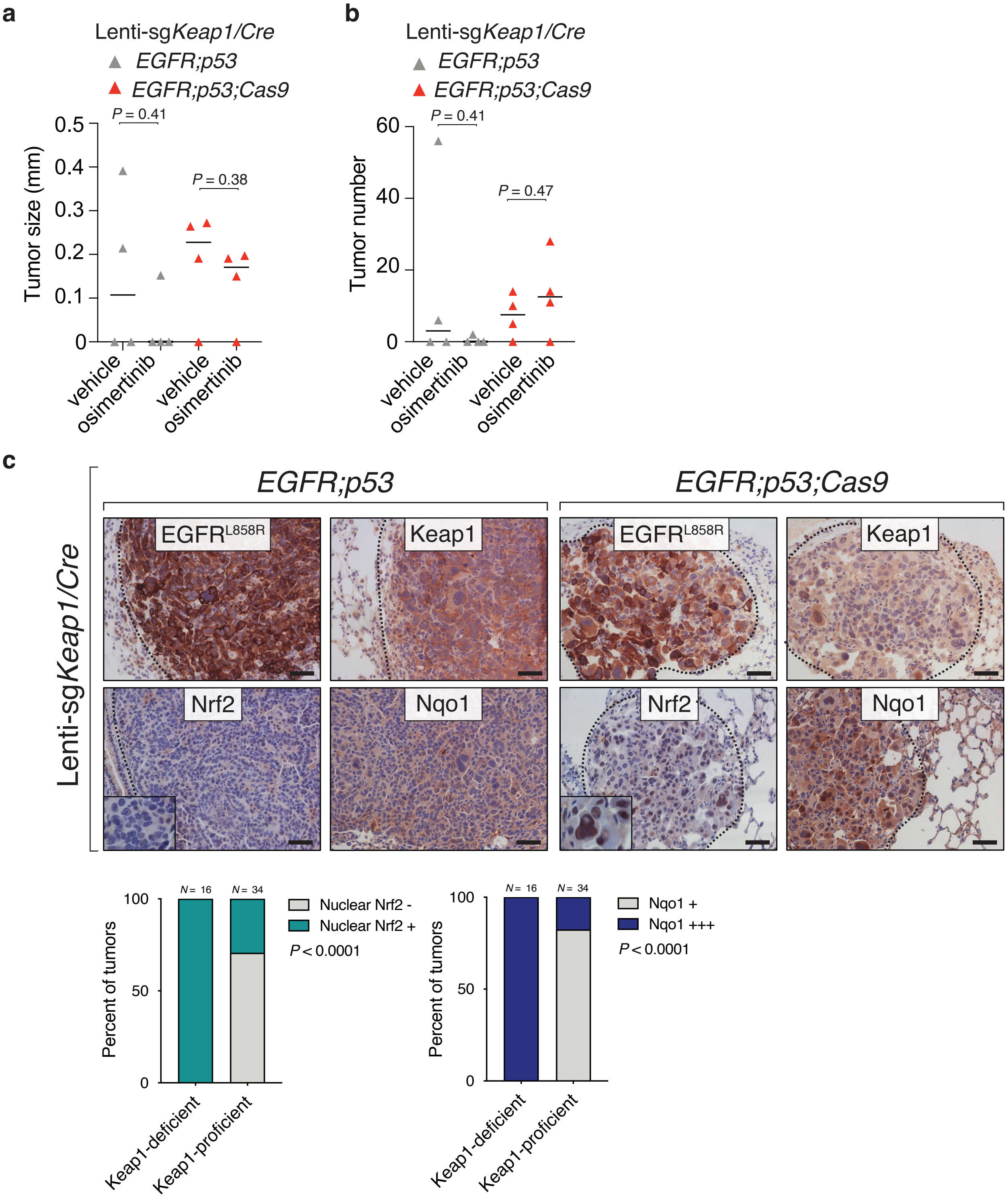


**Supplemental Fig. 7 I Tumor size and number of tumors in vehicle- and osimertinib-treated tumors initiated with individual Lenti-sg*Keap1/Cre* virus. a**, Graph of tumor size in *EGFR;p53* and *EGFR;p53;Cas9* mice with Lenti-sg*Keap1/Cre*-initiated tumors treated with either vehicle or osimertinib for two weeks ($N = 4$ mice-treated/group). Tumor size was calculated by measuring the longest diameter of each tumor in a histological lung section for each group. Each triangle represents the average of all the tumor sizes per mouse. *P*-values were calculated using the Mann-Whitney U test. Horizontal lines show the median. **b**, Graph of tumor number in *EGFR;p53* and *EGFR;p53;Cas9* mice transduced with Lenti-sg*Keap1/Cre* virus and treated with either vehicle or osimertinib for two weeks. In both models, due to large mouse-to-mouse variations, tumor size and number comparisons between vehicle- and osimertinib-treated mice were not significantly different (Mann-Whitney U test). Each triangle represents the number of tumors in one lung section per mouse. Horizontal bars represent the median. **c**, Immunostaining for $EGFR^{L858R}$, Keap1, Nrf2 and Nqo1 in *EGFR;p53* and *EGFR;p53;Cas9* mice with Lenti-sg*Keap1/Cre*-initiated tumors (top left and right panels, respectively). The dashed lines indicate areas of tumors. Scale bars = 50 µm. The panels on the bottom left of the IHC images show nuclear localization of Nrf2 in Keap1-deficient versus Keap1-proficient tumors at higher magnification. Quantification of Nrf2 nuclear expression and Nqo1 expression levels in Keap1-deficient versus Keap1-proficient tumors arising in *EGFR;p53;Cas9* mice (bottom panels). *P*-values were calculated using a Chi-squared test.

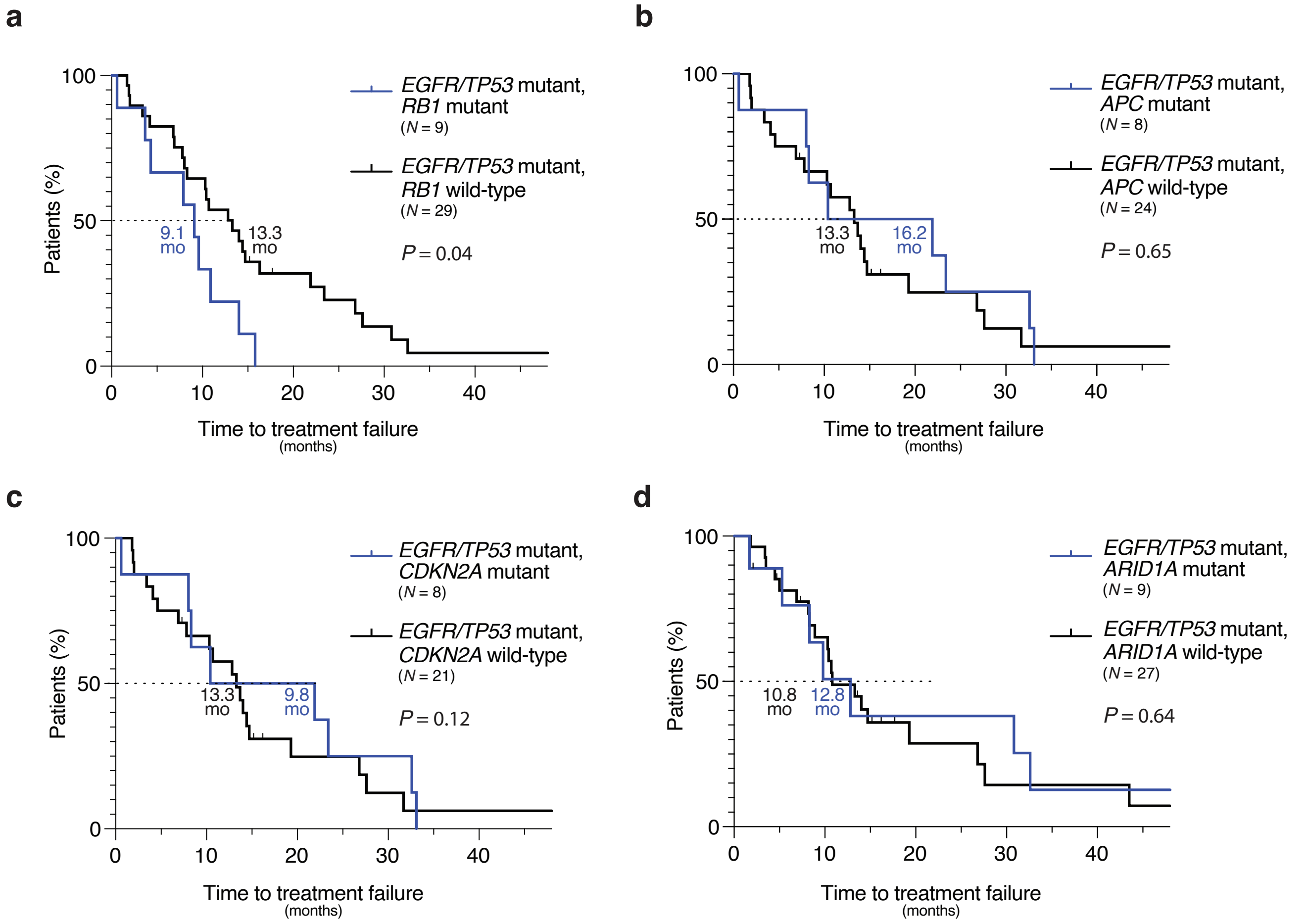


**e**

| Source | *KEAP1* | *NFE2L2* | *CUL3* | TKI therapy | Reference |
|---|---|---|---|---|---|
| human data | loss (1/137) | n.d. | n.d. | AR to Erlotinib | Yu et *al.,* 2018 |
| human data | n.a. | p.W24C (1/21) | n.a. | PD to Erlotinib/ NR to 2nd line Osimertinib | Blakely *et al.,* 2017 |
| human data | n.d. | p.D321N (1/204) | p.N138S (1/204) | AR to Erlotinib | Jordan *et al.,* 2017 |
| *in vitro* | loss | n.d. | n.d. | AR to Erlotinib | Krall *et al.,* 2017 |
| *in vitro* | homozygous mutation (p.D236H) | n.d. | n.d. | AR to Gefitinib | Park *et al.,* 2018 |

**Supplemental Fig. 8 l Mutations in tumor suppressor genes and time to TKI treatment failure in human *EGFR*-driven lung adenocarcinomas. a-d,** Kaplan-Meier curves showing time to TKI treatment failure in human mutant *EGFR/TP53* with mutations in *RB1* (**a**), *APC* (**b**), *CDKN2A* (**c**), *ARID1A* (**d**) compared to mutant *EGFR/TP53* cases without the corresponding genetic alterations. Time to treatment failure was determined by subtracting the date of discontinuation of TKI due to progression, toxicity or death, from the date of initiation of TKI. *P*-values were calcuated using the Log rank test. **e**, Summary of existing literature on *KEAP1/NFE2L2/CUL3* alterations in human mutant *EGFR* lung adenocarcinomas, associated with progressive disease (PD), no response (NR), or acquired resistance (AR) to TKIs. *KEAP1* loss or missense mutations, and missense mutations in *NFE2L2* or *CUL3* have been observed (classified as pathogenic by COSMIC). For each reference, the number of cases, the type of alteration and the TKI therapeutic agent studied are indicated. n.a. = not analyzed. n.d. = not detected.

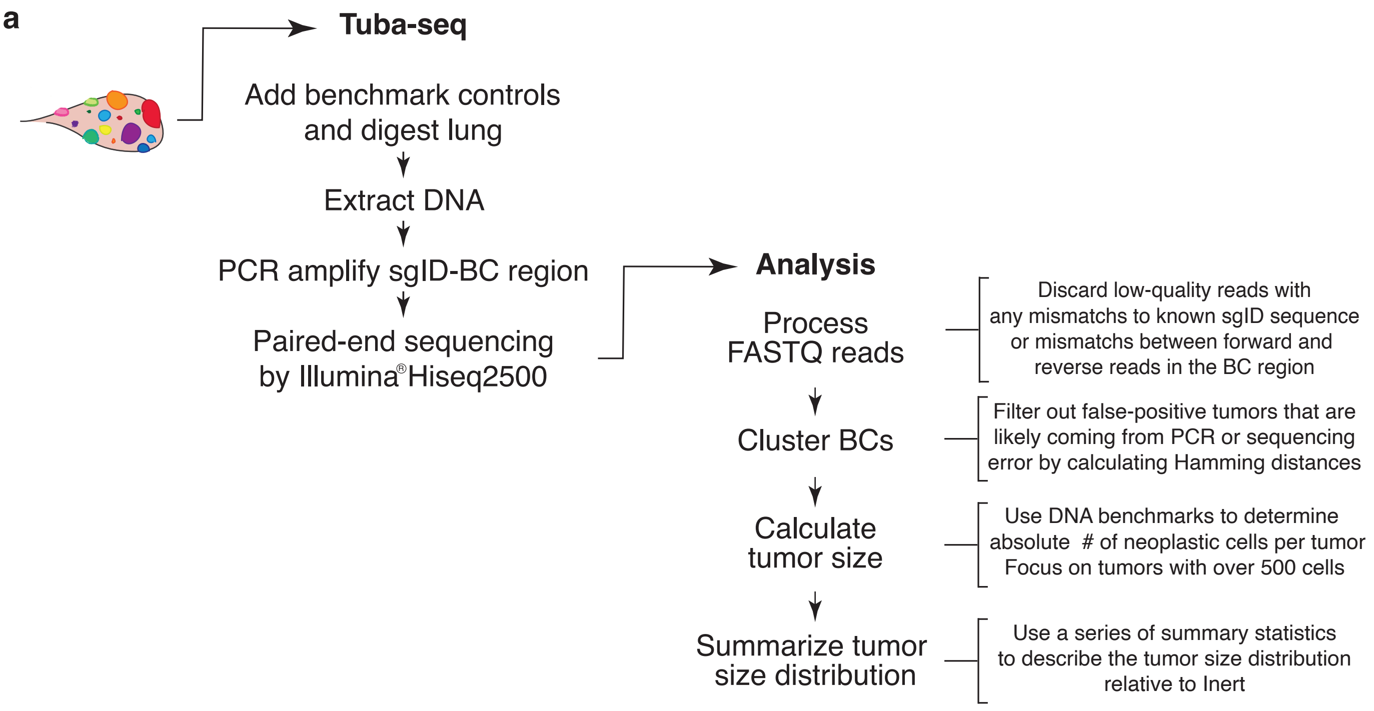


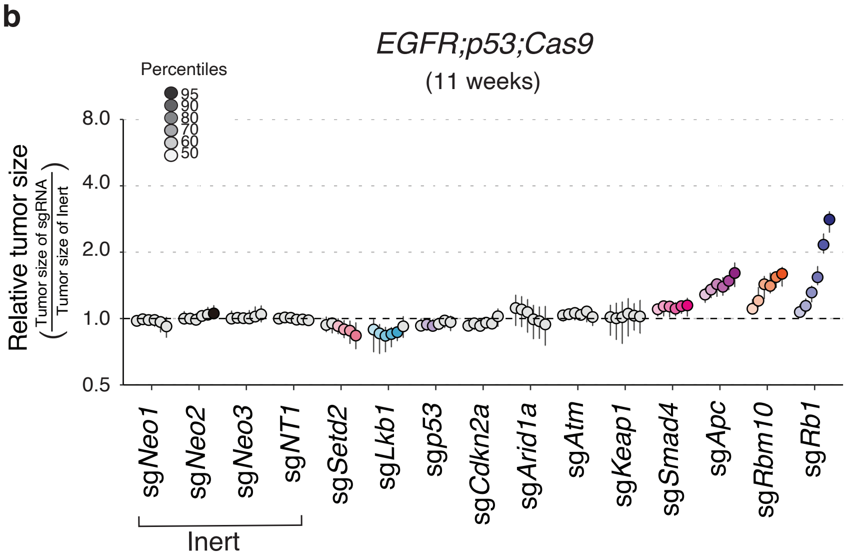


**c**

*EGFR;p53;Cas9*

| | | mean | *P*-value |
|---|---|---|---|
| | sg*Rb1* | **2.17** | **<10$^{-4}$** |
| | sg*Rbm10* | **1.56** | **<10$^{-4}$** |
| | sg*Apc* | **1.53** | **<10$^{-4}$** |
| | sg*Smad4* | **1.15** | **0.02** |
| | sg*Keap1* | 1.03 | 0.77 |
| | sg*Atm* | 1.06 | 0.41 |
| | sg*Arid1a* | 1.05 | 0.66 |
| | sg*Cdkn2a* | 0.97 | 0.22 |
| | sg*p53* | 0.97 | 0.25 |
| | sg*Lkb1* | **0.90** | **<10$^{-4}$** |
| | sg*Setd2* | **0.90** | **0.02** |
| Inert | sg*Neo1* | 0.97 | 0.15 |
| Inert | sg*Neo2* | 1.04 | 0.11 |
| Inert | sg*Neo3* | 1.01 | 0.59 |
| Inert | sg*NT* | 0.99 | 0.34 |

**Supplemental Fig. 9 l Description of the Tuba-seq pipeline analysis and robustness of our model. a**, Tuba-Seq pipeline to quantify tumor sizes *in vivo*. Illumina® sequencing of the DNA barcode region of the integrated lentiviral vectors enables precise measurement of tumor size (see Methods). First, reads with poor quality were discarded. Next, reads were piled-up into groups with unique barcodes. Read pileups were translated into absolute neoplastic cell number using the benchmark controls and by establishing a minimum cell number cutoff to call tumors (cutoff = 500 cells). Lastly, statistical analyses to describe the tumor size distribution were applied. **b**, Relative size distribution of *EGFR;p53;Cas9* tumors at different percentiles 11 weeks after tumor initiation, after simulating a 50% sgRNA efficiency reduction ($N$ = 10). Error bars showing the 95% confidence intervals after bootstrapping. Percentiles that are significantly different from the tumors with inert sgRNAs are in color. **c**, LN mean for tumors with each sgRNA in *EGFR;p53;Cas9* mice 11 weeks after tumor initiation after simulating a 50% sgRNA efficiency reduction (normalized to the tumors with inert sgRNAs). *P*-values are calculated from bootstrapping. *P*-values < 0.05 and their corresponding means are highlighted for sgRNAs that positively (red) and negatively (green) affect tumor growth when the effects are equal to or differ >10% compared to the size of tumors with inert sgRNAs (Right panel).

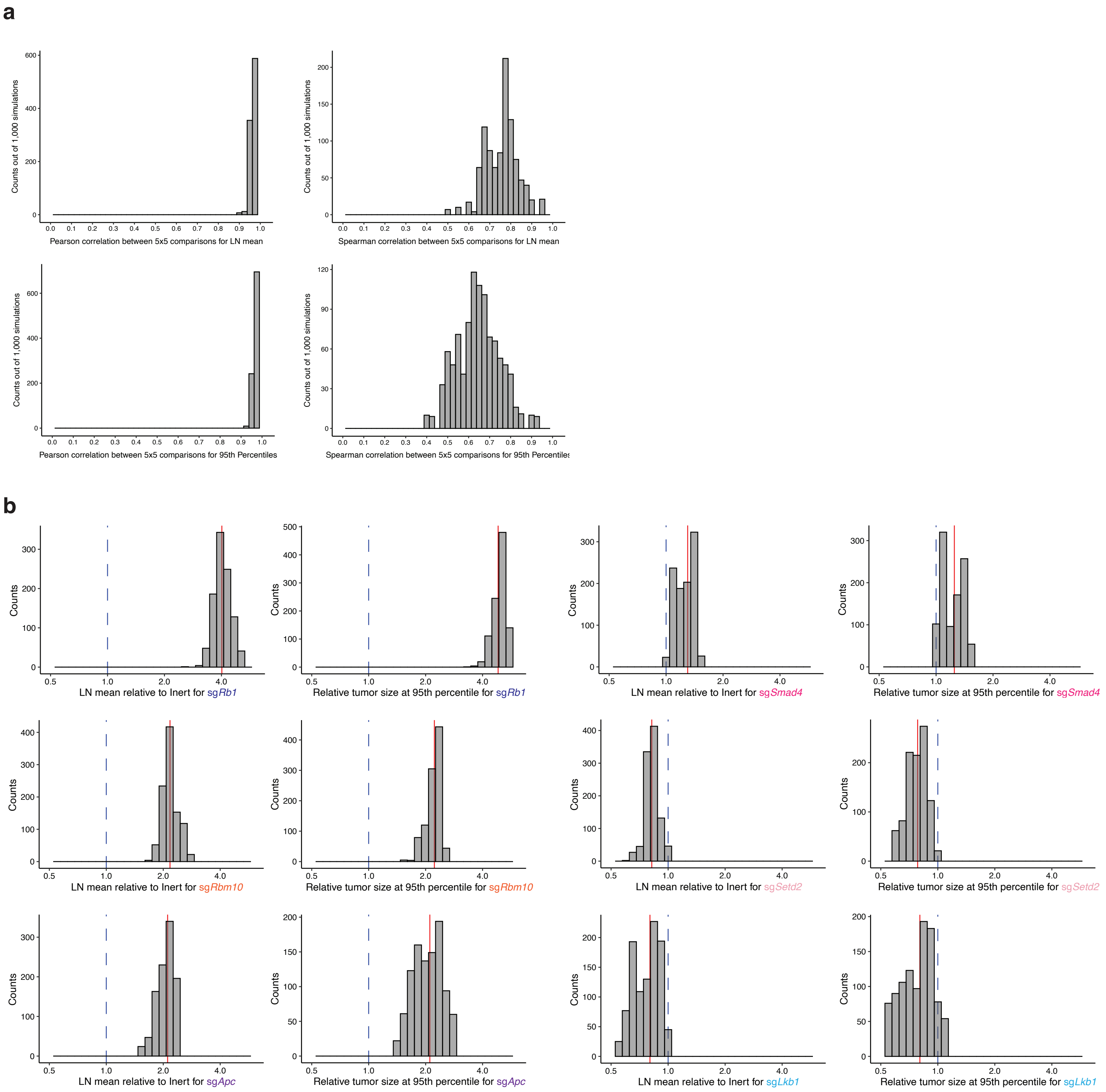


**Supplemental Fig. 10 l Replicate reproducibility supports Tuba-seq results. a,** Pearson and Spearman correlation between LN mean and 95th percentile of tumor size distribution at 11 weeks of tumor initiation after performing subsampling into two groups of 5 mice (*N* = 10). **b,** Relative LN mean and 95$^{th}$ percentile distribution of tumor size (left and right panels for each sg*TS*, respectively) obtained by reducing the number of mice by half (5 mice/group). The dotted blue line indicates no effect, and the solid red line is the estimated value of LN mean or the 95$^{th}$ percentile distribution using all 10 mice at 11 weeks of tumor initiation. This analysis confirmed our results on the functional effects of these sgRNAs in *EGFR;p53;Cas9* mice. Reducing the number of mouse by half still yields quantitatively similar results supporting that our sample size is sufficiently large.

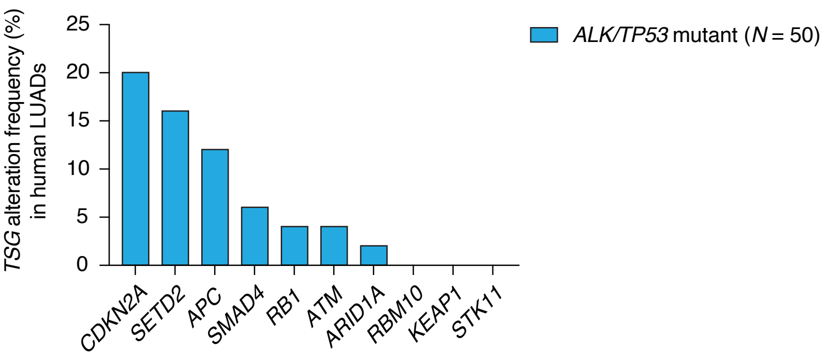


**Supplemental Fig. 11 | Tumor suppressor gene mutational spectrum in lung adenocarcinomas with *ALK* rearrangements and *TP53* mutations.** Frequency of tumor suppressor gene alterations that co-occur with *ALK* rearrangements and *TP53* mutations in human lung adenocarcinomas (LUADs, data from AACR Project GENIE). Each bar represents the alteration frequency for each tumor suppressor gene in human lung adenocarcinomas. Interestingly, *ALK*-driven tumors show a strikingly different mutation spectrum for tumor suppressor genes, with *SETD2, CDKN2A,* and *APC* being the most commonly co-occurring alterations in this setting, compared to *EGFR*-driven tumors (Fig. 4d). Notably, *RBM10* constrains both *EGFR*- and *KRAS*-driven tumors *in vivo* and it is one of the most frequently altered genes in both oncogenic contexts, but it is not found altered in *ALK*-driven lung tumors in this dataset.

| **Characteristics** | ***KEAP1/NFE2L2/CUL3*** **mutant (n = 10)** | ***KEAP1/NFE2L2/CUL3*** **wild-type (n = 30)** | ***P*-value** |
|---|---|---|---|
| Age, mean (years) | 64 | 61 | 0.77 |
| Sex | | | |
| Male | 3 | 11 | 1.0 |
| Female | 7 | 19 | |
| Race | | | |
| White | 4 | 13 | 1.0 |
| Asian | 6 | 17 | |
| Smoking History | | | |
| Current | 0 | 0 | |
| Previous | 3 | 10 | 0.1 |
| Never | 7 | 20 | |
| EGFR mutation | | | |
| L858R | 6 | 11 | |
| Exon 19 deletion | 2 | 18 | 0.11 |
| Other* | 2 | 1 | |
| TKI Therapy | | | |
| Erlotinib | 7 | 26 | |
| Osimertinib | 2 | 3 | 0.57 |
| Other+ | 1 | 1 | |

*Other: exon 21 mutation, exon 20 insertion. +: afatinib, TKI on clinical trial.

**Supplemental Table 1 I NSCLC cohort generation from the Stanford Solid Tumor Actionable Mutation Panel (STAMP).** Demographic data from the STAMP cases of patients with stage IV non-small cell lung cancer (NSCLC), oncogenic *EGFR* and *TP53* mutations included in the study. Data including sex, age at diagnosis, race, smoking history, type of *EGFR* mutation and TKI therapy are reported. The cohort was matched to a *KEAP1/NFE2L2/CUL3* wild-type cohort and no significant difference was observed based on the matched variables.

| | Hazard Ratio | 95% CI | *P*-value |
|---|---|---|---|
| *KEAP1* mutation | 2.6 | 1.05-6.43 | 0.04* |
| Age at diagnosis | 1.0 | 0.97-1.03 | 0.99 |
| Sex | 1.5 | 0.79-2.65 | 0.23 |
| Tobacco Use | 1.3 | 0.54-3.11 | 0.57 |
| Race | 0.9 | 0.41-1.77 | 0.67 |

**Supplemental Table 2 I Impact of confounding variables on time to treatment failure with TKIs.** Even after adjustment for potential confounders such as age, sex, race and smoking status, the multivariate analysis shows that *KEAP1* mutations significantly reduced time on treatment with TKIs.

| **Characteristics** | ***EGFR* mutant (n = 29)** |
|---|---|
| Age, mean (years) | 60 |
| Sex | |
| Male | 11 |
| Female | 18 |
| Race | |
| Black | 2 |
| White | 25 |
| Asian | 2 |
| Smoking History | |
| Current | 0 |
| Previous | 16 |
| Never | 13 |
| EGFR mutation | |
| L858R | 12 |
| Exon 19 deletion | 15 |
| G719X | 2 |
| TKI Therapy | |
| Erlotinib | 26 |
| Gefitinib | 1 |
| Afatinib/Cetuximab | 2 |

**Supplemental Table 3 I NSCLC cohort from the Yale Lung Rebiopsy Program (YLR).** Demographic data from the available YLR cases of patients with stage IV NSCLC and mutant *EGFR* included in the study. Data including sex, age at diagnosis, race, smoking history, type of *EGFR* mutation and TKI therapy are reported.

| Vectors for Lenti-sgTS$^{Pool}$/Cre | sgRNA Sequence | Target Exon [Total # of Exons] | Adjacent to SA/SD | Upstream of Functional Domain [Functional Domain] | References |
|---|---|---|---|---|---|
| Lenti-sgApc/Cre | TTGAGCGTAGTTTCACTCCG | Exon 4 [16] | No | Yes [Eb1] | Rogers *et al.,* 2017 |
| Lenti-sgArid1a/Cre | TATGGGTTAGTCCCACCATA | Exon 2 [20] | No | Yes [ARID] | Rogers *et al.,* 2017 |
| Lenti-sgAtm/Cre | GCTAAGATGTGACTTAAGCC | Exon 7 [64] | No | Yes [FAT, PI3K/PI4K, FATC] | Rogers *et al.,* 2017 |
| Lenti-sgCdkn2a/Cre | GCGCTGCGTCGTGCACCGGG | Exon 2 [3] | No | N/A | Rogers *et al.,* 2017 |
| Lenti-sgKeap1/Cre | CCTGCACGTGATGAACGGGG | Exon 2 [6] | No | Yes [BACK] | Rogers *et al.,* 2017 |
| Lenti-sgLkb1/Cre | GTGGTGGGCCGCAGTCACAA | Exon 6 [10] | No | Yes [Protein Kinase] | Rogers *et al.,* 2017 |
| Lenti-sgp53/Cre | AGGAGCTCCTGACACTCGGA | Exon 4 [8] | No | Yes [DNA-binding, Tetramer] | Rogers *et al.,* 2017 |
| Lenti-sgRb1/Cre | TCTTACCAGGATTCCATCCA | Exon 2 [26] | No | N/A | Rogers *et al.,* 2017 |
| Lenti-sgRbm10/Cre | GTATTTCCTGAACAGATCCG | Exon 5 [24] | Yes | Yes [RRM1, RRM2, G-Patch] | Rogers *et al.,* 2017 |
| Lenti-sgSetd2/Cre | TCTCTAATCCATCTTCCCAG | Exon 3 [21] | No | Yes [AWS, SER, Post-SET, WW] | Rogers *et al.,* 2017 |
| Lenti-sgSmad4/Cre | GGTGGCGTTAGACTCTGCCG | Exon 5 [12] | No | Yes [MH2] | Rogers *et al.,* 2017 |
| Lenti-sgNeo1/Cre | TCATGGCTGATGCAATGCGG | N/A | N/A | N/A | Rogers *et al.,* 2017 |
| Lenti-sgNeo2/Cre | GATATTGCTGAAGAGCTTGG | N/A | N/A | N/A | Rogers *et al.,* 2017 |
| Lenti-sgNeo3/Cre | GAATAGCCTCTCCACCCAAG | N/A | N/A | N/A | Rogers *et al.,* 2017 |
| Lenti-sgNT/Cre | GCGAGGTATTCGGCTCCGCG | N/A | N/A | N/A | Rogers *et al.,* 2017 |

**Supplemental Table 4 I List of selected sgRNAs targeting candidate tumor suppressor genes.** Vectors designed to initiate tumors with Lenti-*TS$^{Pool}$/Cre* virus. All vectors have been published in Rogers *et al.,* 2017.

| Vectors for Lenti-sgTS/Cre | sgRNA Sequence | Target Exon [Total # of Exons] | Adjacent to SA/SD | Upstream of Functional Domain [Functional Domain] | References |
|---|---|---|---|---|---|
| Lenti-sg*Apc/Cre* | TTGAGCGTAGTTTCACTCCG | Exon 4 [16] | No | Yes [Eb1] | Rogers *et al.,* 2017 |
| Lenti-sg*Rbm10/Cre* | GTATTTCCTGAACAGATCCG | Exon 5 [24] | Yes | Yes [RRM1, RRM2, G-Patch] | Rogers *et al.,* 2017 |
| Lenti-sg*Rbm10#2/Cre* | CTGTCTCCAGGTCAGAGCCG | Exon 6 [24] | No | Yes [RRM1, RRM2, G-Patch] | |
| Lenti-sg*Keap1/Cre* | CCTGCACGTGATGAACGGGG | Exon 2 [6] | No | Yes [BACK] | Rogers *et al.,* 2017 |
| Lenti-sg*Neo2/Cre* | GATATTGCTGAAGAGCTTGG | N/A | N/A | N/A | Rogers *et al.,* 2017 |

**Supplemental Table 5 I List of selected sgRNAs to validate the role of tumor suppressor gene inactivation on tumor growth and therapeutic sensitivity.** Vectors designed to initiate tumors with individual Lenti-sg*TS/Cr*e virus.
